# Cryo-EM structures of cardiac thin filaments reveal the 3D architecture of troponin

**DOI:** 10.1101/2020.01.15.908418

**Authors:** Toshiyuki Oda, Haruaki Yanagisawa, Takeyuki Wakabayashi

**Affiliations:** Department of Anatomy and Structural Biology, Graduate School of Medicine, University of Yamanashi, 1110 Shimokato, Chuo, Yamanashi, 409-3898, Japan.; Department of Cell Biology and Anatomy, Graduate School of Medicine, the University of Tokyo, 7-3-1 Hongo, Bunkyo-ku, Tokyo, 113-0033, Japan.; The University of Tokyo, 7-3-1 Hongo, Bunkyo-ku, Tokyo, 113-0033, Japan.

**Keywords:** Cryo-electron microscopy, Cardiac muscle, Actin, Troponin, Tropomyosin

## Abstract

Troponin is an essential component of striated muscle and it regulates the sliding of actomyosin system in a calcium-dependent manner. Despite its importance, the structure of troponin has been elusive due to its high structural heterogeneity. In this study, we analyzed the 3D structures of murine cardiac thin filaments using a cryo-electron microscope equipped with a Volta phase plate (VPP). Contrast enhancement by a VPP enabled us to reconstruct the entire repeat of the thin filament. We determined the orientation of troponin relative to F-actin and tropomyosin, and characterized the interactions between troponin and tropomyosin. This study provides a structural basis for understanding the molecular mechanism of actomyosin system.

## Introduction

Contraction of striated muscles is driven by the interaction between the thin (actin) and the thick (myosin) filaments. The main components of thin filaments are F-actin, tropomyosin, and troponin. Tropomyosin molecules assumes a coiled-coil structure and bind to F-actin. Together with troponin, tropomyosin blocks actin-myosin binding under low calcium condition (Behrmann et al., 2012; von der Ecken et al., 2015). When intracellular calcium concentration is elevated, calcium-binding to troponin induces a shift in tropomyosin to expose the myosin-binding sites of F-actin, allowing muscle contraction (Gordon et al., 2000).

Troponin is a complex composed of three subunits: troponin C (TnC), troponin I (TnI), and troponin T (TnT) (Greaser and Gergely, 1971; Ebashi et al., 1971). TnC is composed of N-terminal and C-terminal globular domains. Calcium-binding to the N-terminal domain triggers the muscle contraction (Strynadka et al., 1997). TnI has an actin-binding inhibitory region, which is essential for the inhibition of actin-myosin interaction (Ramos, 1999). TnT binds to tropomyosin and anchors the troponin complex to the thin filament (Ebashi et al., 1971; Mak and Smillie, 1981).

Structural analysis of troponin, especially with cryo-EM, has been hindered by its sparse localization, low contrast, and high structural heterogeneity. Therefore, 3D structural analyses of troponin have been relying on negative-staining method (Yang et al 2014; Paul et al., 2017). Although heavy-metal staining increases the contrast of troponin, resolution of the negatively-stained troponin map (∼27 Å) is insufficient to make a meaningful comparison with crystal/NMR structures. To enhance the contrast of frozen-hydrated troponin and improve the accuracy of alignment, we utilized the VPP technology (Danev and Baumeister, 2016) for the cryo-EM of cardiac thin filaments. Focused 3D classification and multi-body refinement (Nakane et al., 2018) successfully solved the structures of thin filaments in sub-nanometer resolution and revealed calcium-dependent changes in the structural relationship between troponin and tropomyosin. This study has uncovered the long-sought piece of the puzzle in understanding the regulatory mechanism of actomyosin system.

## Results

### Cryo-EM of cardiac thin filaments

We isolated thin and thick filaments from murine myocardium and reconstructed the 3D structures of the thin filaments in low- and high-calcium states. We initially failed to visualize the troponin structure on native thin filaments probably due to detachment of troponin upon blotting and freezing. Thus, we fixed the filaments using 0.25% glutaraldehyde (Risi et al., 2017), which greatly improved the occupancy of troponin on the filaments (Fig. 1A). Fortunately, it has been reported that glutaraldehyde-fixation does not inhibit the calcium-dependent activation of myosin by thin filaments, although the efficiency of myosin-activation and mobility of tropomyosin were slightly altered (Risi et al., 2017). The effect of fixation is further discussed later (see Discussion).

**Fig. 1.**
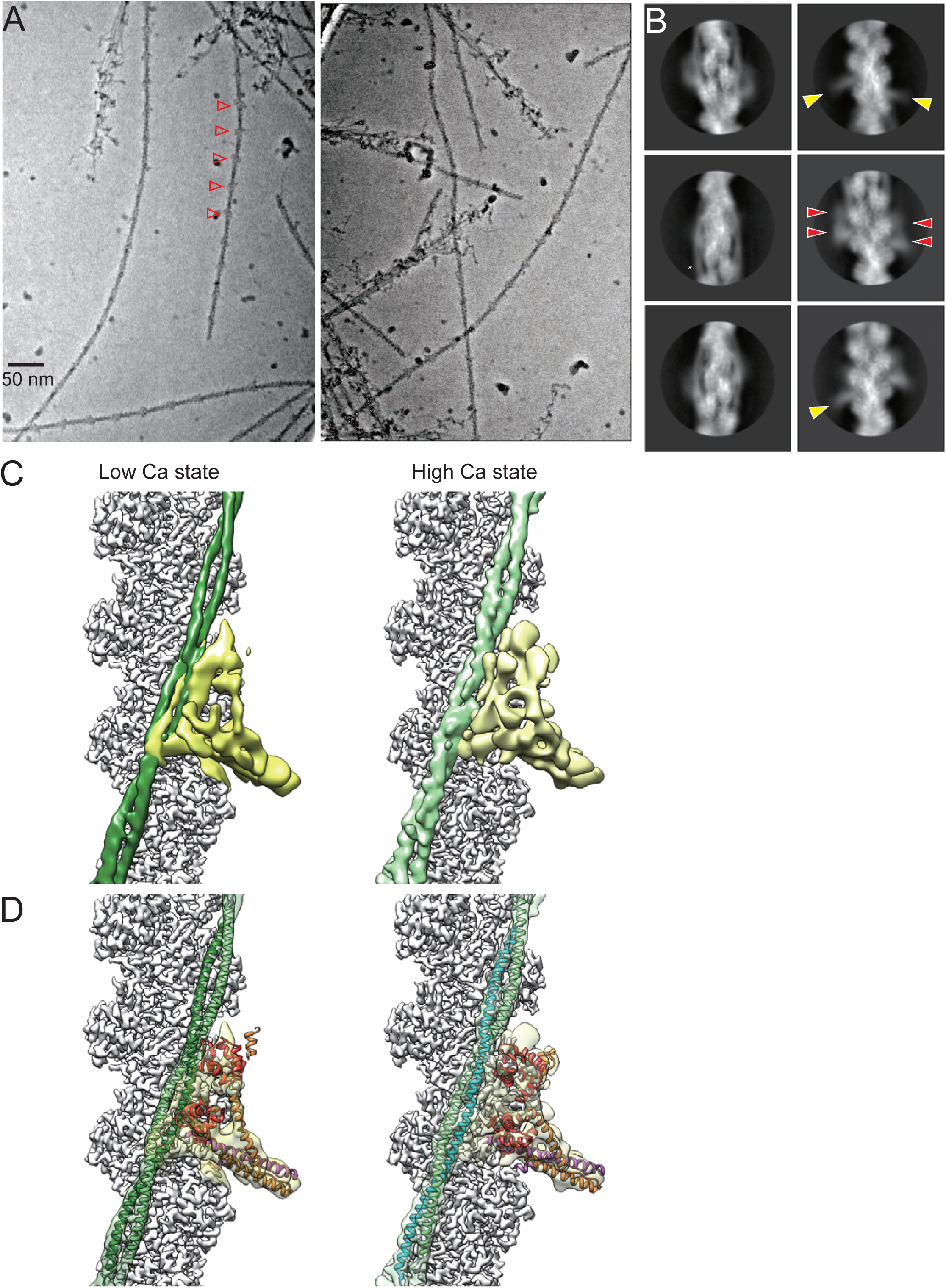
Cryo-electron microscopy of cardiac thin filaments. (A) Electron micrographs of frozen-hydrated cardiac thin filaments. Left, low calcium state; right, high calcium state. Arrowheads indicate the troponin heads. **(B)** Representative 2D class averages of the troponin-containing segments. Red arrowheads indicate the globular TnCN and TnCC domains. Yellow arrowheads indicate the “tail-shaped” IT arms. **(C)** Results of the three-body refinement reassembled into one volume. Gray: F-actin pointed-end up; green: tropomyosin; yellow: troponin. **(D)** Troponin and tropomyosin models were fitted into the density maps. Red: TnC; purple: TnT; orange: TnI; green/cyan: tropomyosin. **(C-D**, right) Enlarged images to show the shifts in tropomyosin between the two calcium states.

Image acquisition with a VPP significantly increased the contrast (Fig. 1A-B). However, troponin densities were dispersed in the reconstructed maps of the thin filaments (Fig. S1-2, upper right). To improve the alignment of troponin, we subtracted actin densities from the original images (Roh et al., 2017), and conducted a 3D classification focusing on the troponin region (Zivanov et al., 2018). Although 3D classification could refine the troponin region (Fig. S1-2), densities of actin and tropomyosin outside the troponin mask were disordered, due to the variation in the location of troponin relative to actin and tropomyosin. Therefore, we performed multi-body 3D refinements (Nakane et al., 2018). The refined maps clearly revealed the 3D organization of the entire thin filament complex, as well as the calcium-induced shift of tropomyosin (Fig. 1C-D, and Movies S1-2).

### Orientation and domain organization of troponin

The orientation of troponin relative to F-actin and tropomyosin has been controversial due to the insufficient structural information (Paul et al., 2017; Yang et al., 2014; Narita et al., 2001). The most widely accepted model is that the TnI-TnT arm (IT arm) lies parallel to tropomyosin (Fig. 2A-B, right) (Paul et al., 2017). Our structures, however, disagreed with this “inverted” model, because the crystal structure in the “inverted” orientation could not be fitted into our map (Fig. 2A, right). Instead, our structures suggest a new model in that the IT arm lies perpendicular to tropomyosin (Fig. 2A, middle; Fig. 2B, left).

**Fig. 2.**
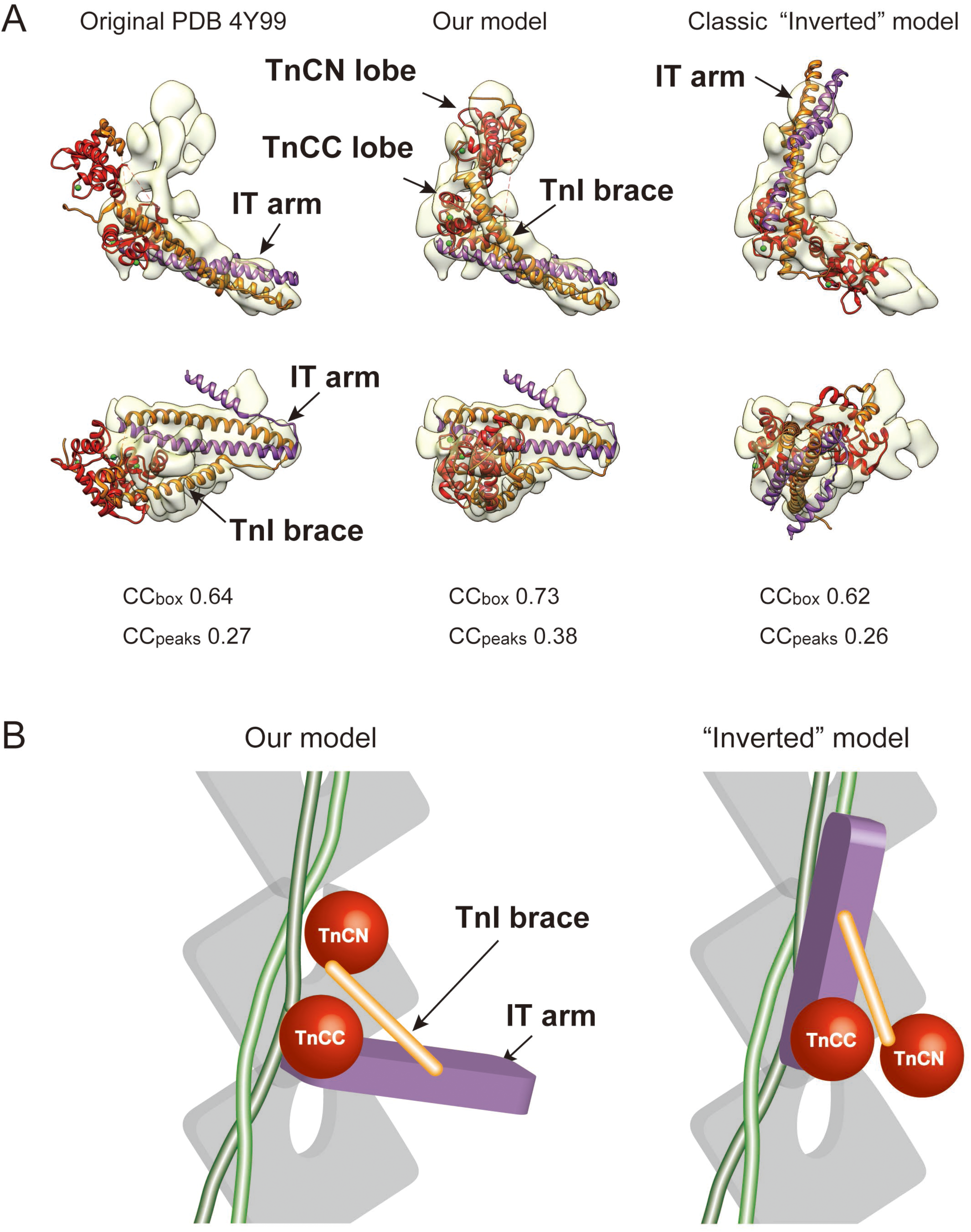
Comparison between our model and previous “inverted” model of troponin. (A) Fitting of the crystal structure PDB 4Y99 into our troponin map. Lower panels are top views from the pointed-end. The PDB 4Y99 model was first manually fitted into our troponin in different orientations using UCSF Chimera. The manually fitted models were further aligned using PHENIX real space refinement. Cross-correlation values calculated using PHENIX comprehensive validation tool were indicated below the maps. When the PDB 4Y99 model was fitted directly into the map, TnC (red) is largely outside of the map due to the “open-up” configuration (left). Rotating the TnC gave a reasonable fit (middle). Fitting was obviously failed when the IT arm was put in parallel with tropomyosin (Classic “inverted” model, right). Purple: TnT; orange: TnI. **(B)** In our model, the TnCN and TnCC lobes (red) lie parallel to tropomyosin (green). In the “inverted” model, the IT arm (purple) lies parallel to tropomyosin. The TnI brace (orange) bridges between the TnC lobes and the IT arm. Gray: actin.

Our cryo-EM structures showed that troponin took an “L”-shape, which were composed of two globular domains and one flat, plate-like domain (Fig. 2B). Comparison with the crystal structure of human cardiac troponin core complex in high-calcium state (PDB 4Y99, Takeda et al., 2003) suggests that the two globular domains correspond to the two lobes of TnC (Dvoretsky et al., 2002), which were designated as “TnCN” and “TnCC” lobes (Fig. 2A-B, red). The plate-like domain, which appeared as a thin tail in the 2D class averages (Fig. 1B, yellow arrowheads), is considered to be the IT arm (Fig. 2B, purple). Interestingly, the crystal structure cannot be fitted directly into our map and appears to take an “open-up” conformation, suggesting presence of a hinge at the corner of the L-shape (Fig. 2A, left). Tilting TnC by ∼50 ° counterclockwise about this hinge gave a reasonable fit onto our map (Fig. 2B, middle). This new model of troponin orientation is consistent with previous biochemical studies (Kimura-Sakiyama et al., 2008; Ferguson et al., 2003, see Discussion for details).

Between the TnCN lobe and the IT arm, there is a diagonal brace, which was designated as “TnI brace” (Fig. 2B, orange). In the crystal structure, the N-terminal alpha-helix of TnI similarly bridges between the TnCN and the IT arm (Fig. 2A), thus the TnI brace is likely to be composed of the N-terminal alpha-helix of TnI.

### Calcium-dependent conformational changes

Comparison between the structures in low- and high-calcium states suggests that the tropomyosin shift is induced by conformational changes in troponin (Fig. 3). In low-calcium state, the IT arm formed a bent at one end (Fig. 3A, arrowhead) and ran between the two alpha-helices of tropomyosin. This domain, designated as “TnT hook”, is probably the C-terminal domain (T2) of TnT, which tightly associates with tropomyosin (Perry, 1998; Katrukha, 2013, see Discussion). In high-calcium state, by contrast, the TnT hook translocated sideways to put itself between TnC and tropomyosin (Fig. 3B, arrowheads), which made an ∼11 Å shift in tropomyosin (Fig. 3A-C, green).

**Fig. 3.**
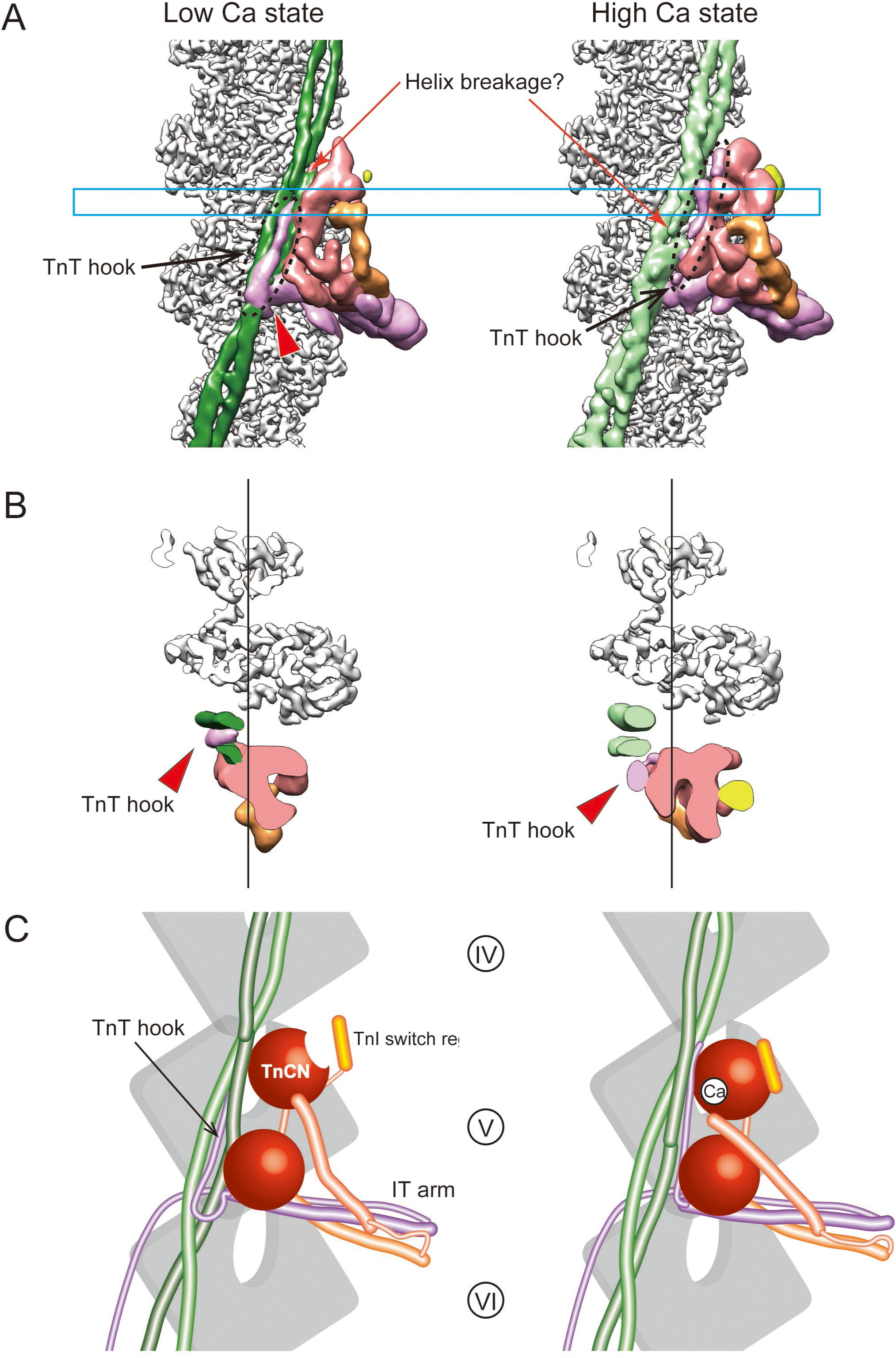
Calcium-dependent structural changes in troponin and tropomyosin. (A) Close-up views of troponin and tropomyosin. Arrowhead indicates the bending in the IT arm, which is connected to the TnT hook. Cyan box indicates the position of the section in B. Red: TnC; orange: TnI brace; yellow: TnI switch region; purple: IT arm. **(B)** Cross-sectional views. Vertical lines were placed to clearly show the tropomyosin shift. **(C)** Schematic of the calcium-dependent conformational changes. (A, C) In low-calcium state, the TnCN lobe appeared thinner than that in high-calcium state, presumably because of the calcium-dependent binding of the switch region of TnI (yellow) to TnCN lobe (Gordon et al., 2000). Note that the C-terminal 16 residues of TnT, corresponding to the TnT hook, is not visualized in the crystal structure (Takeda et al 2003).

Interestingly, there was an apparent breakage in one of the helices of tropomyosin in both calcium states (Fig. 3A, orange arrows). This structural feature may facilitate the calcium-dependent displacement of tropomyosin, but further studies are necessary to understand its physiological meaning.

To investigate the effect of tropomyosin shift on actin-myosin binding, we reconstructed the entire repeat of the thin filament, which includes the whole 385 Å-long tropomyosin coiled-coil dimer (Fig. S3A). We examined each seven actin level (Fig. 4A-G, Table 1, Fig. S3B, S4A) by fitting an actin-myosin model (gray and deep blue, Mentes et al., 2018) and actin-tropomyosin models (red, Risi et al 2017). Note that the “helix 1” was defined as the alpha-helix that faces troponin at the level V. Because the tropomyosin colied-coil makes one turn at each actin level, the helix 1 and 2 interfere with the myosin head at the odd-numbered and the even-numbered levels, respectively.

**Fig. 4.**
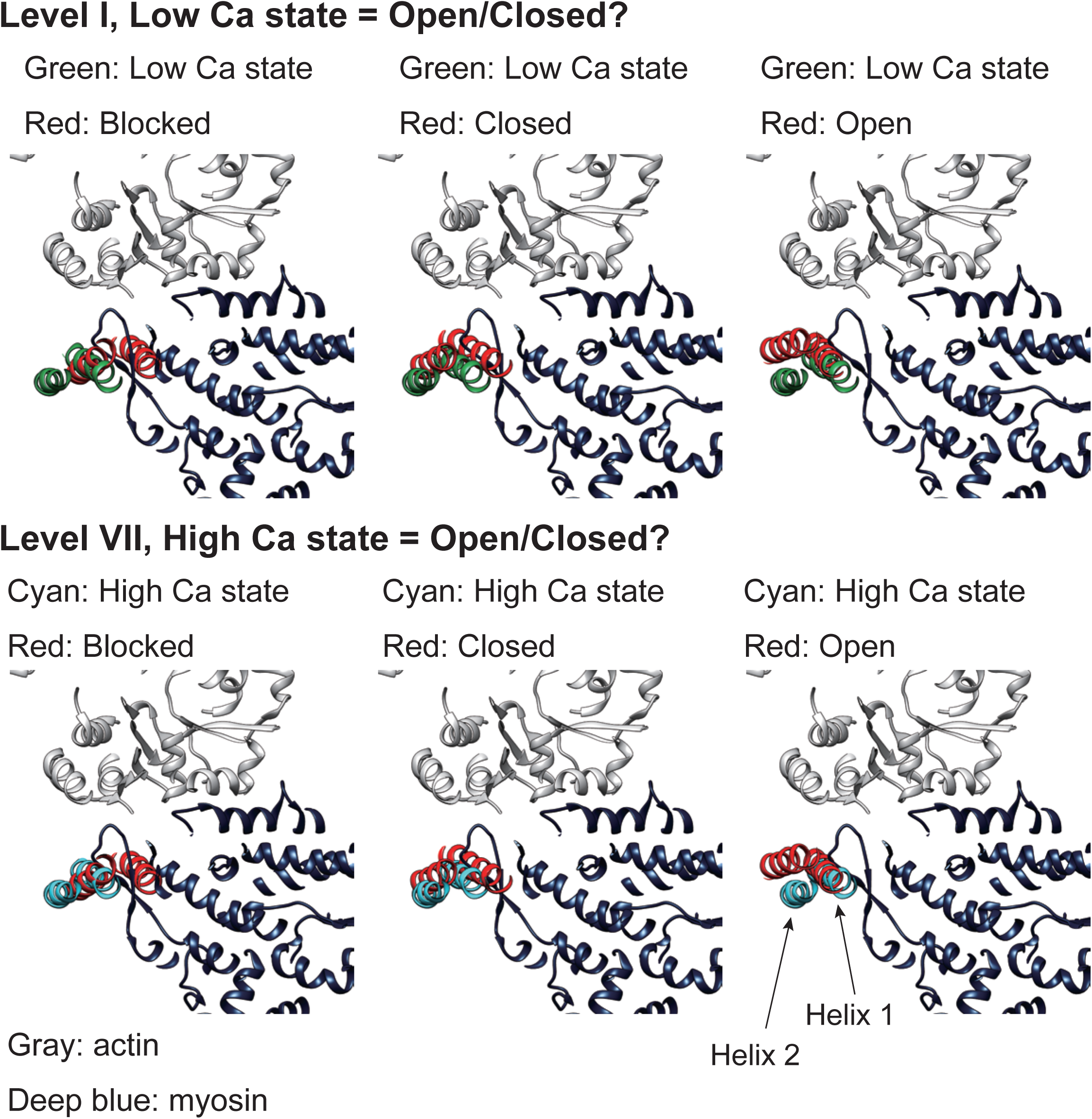
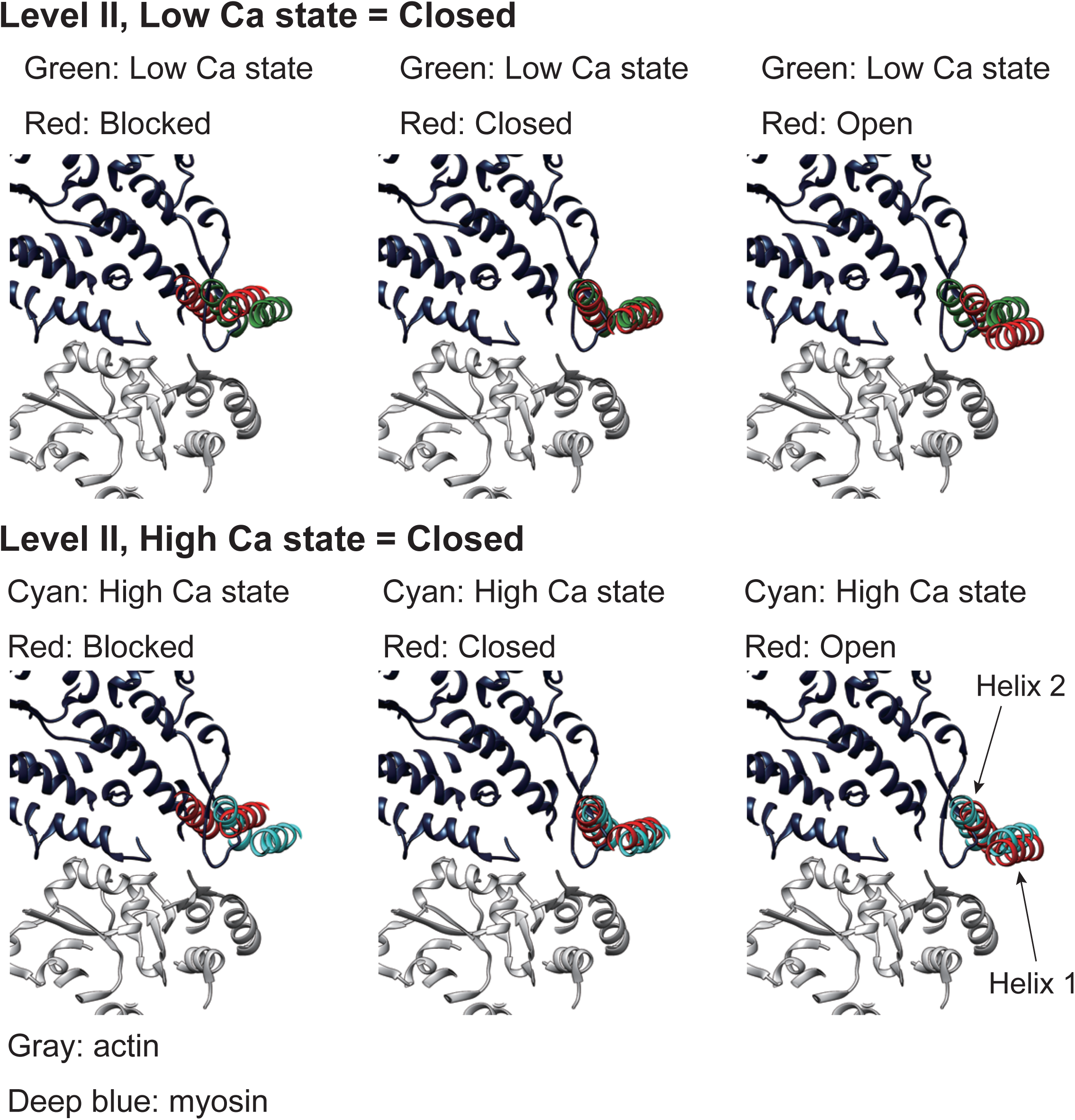
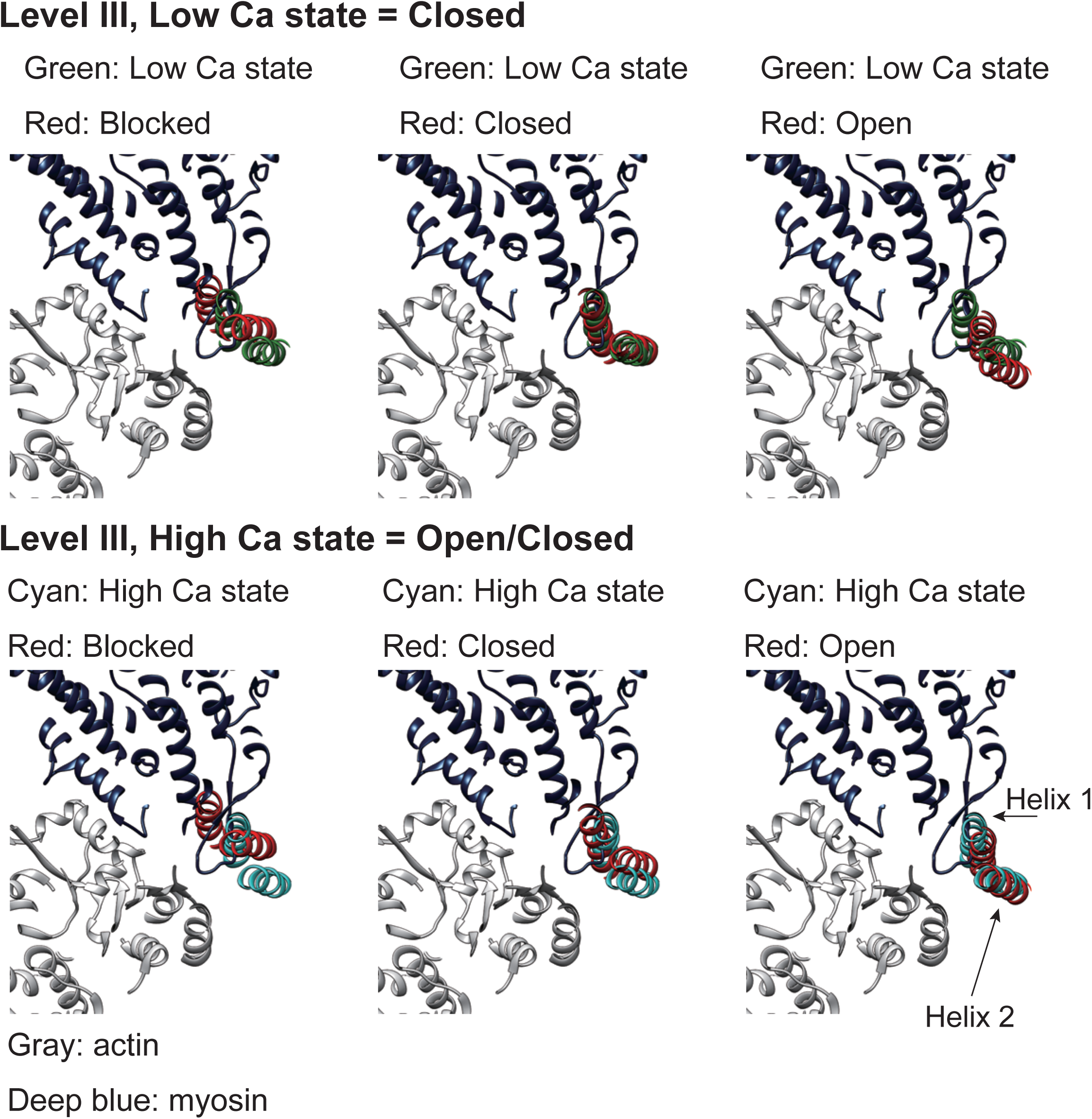
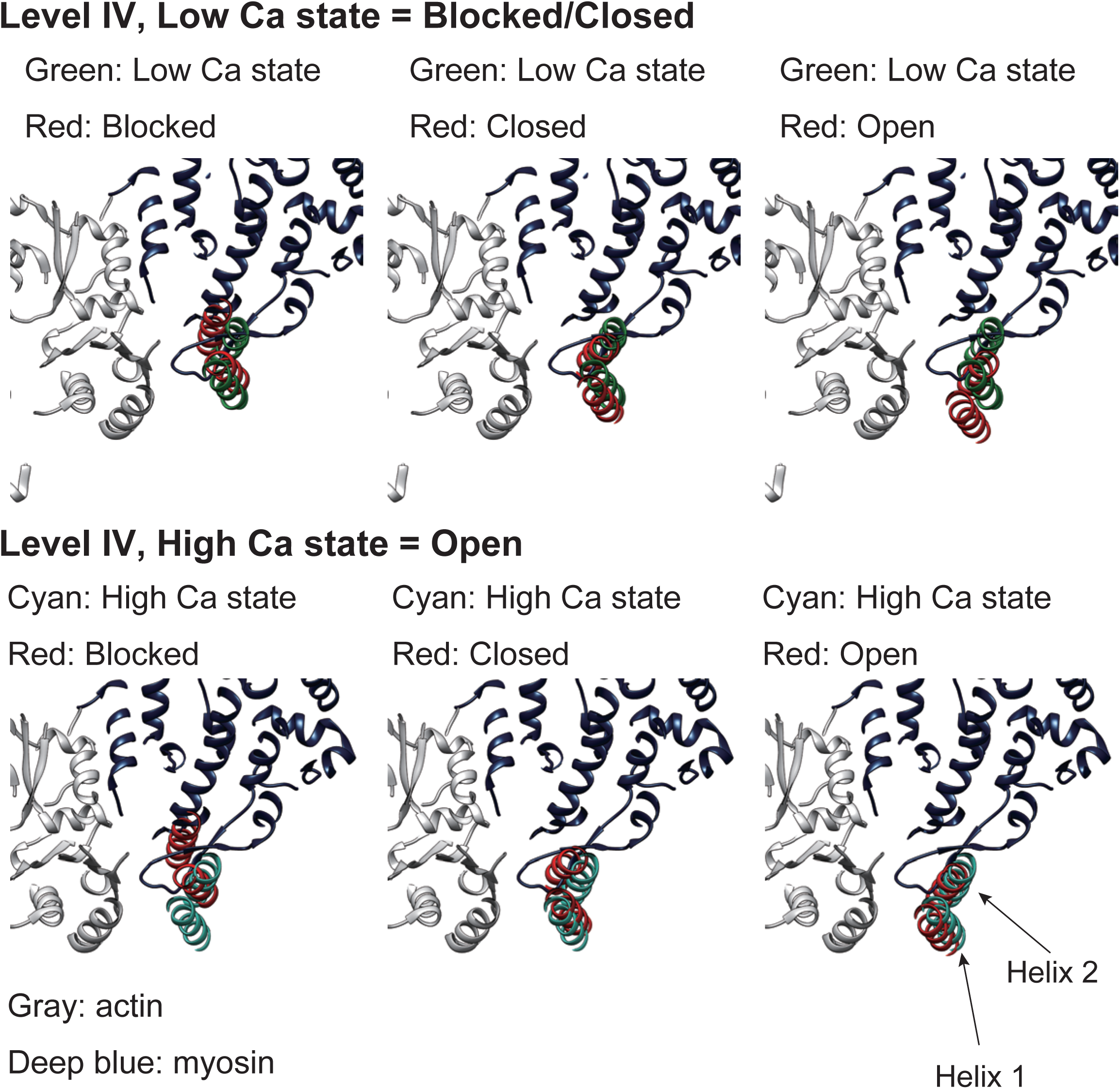
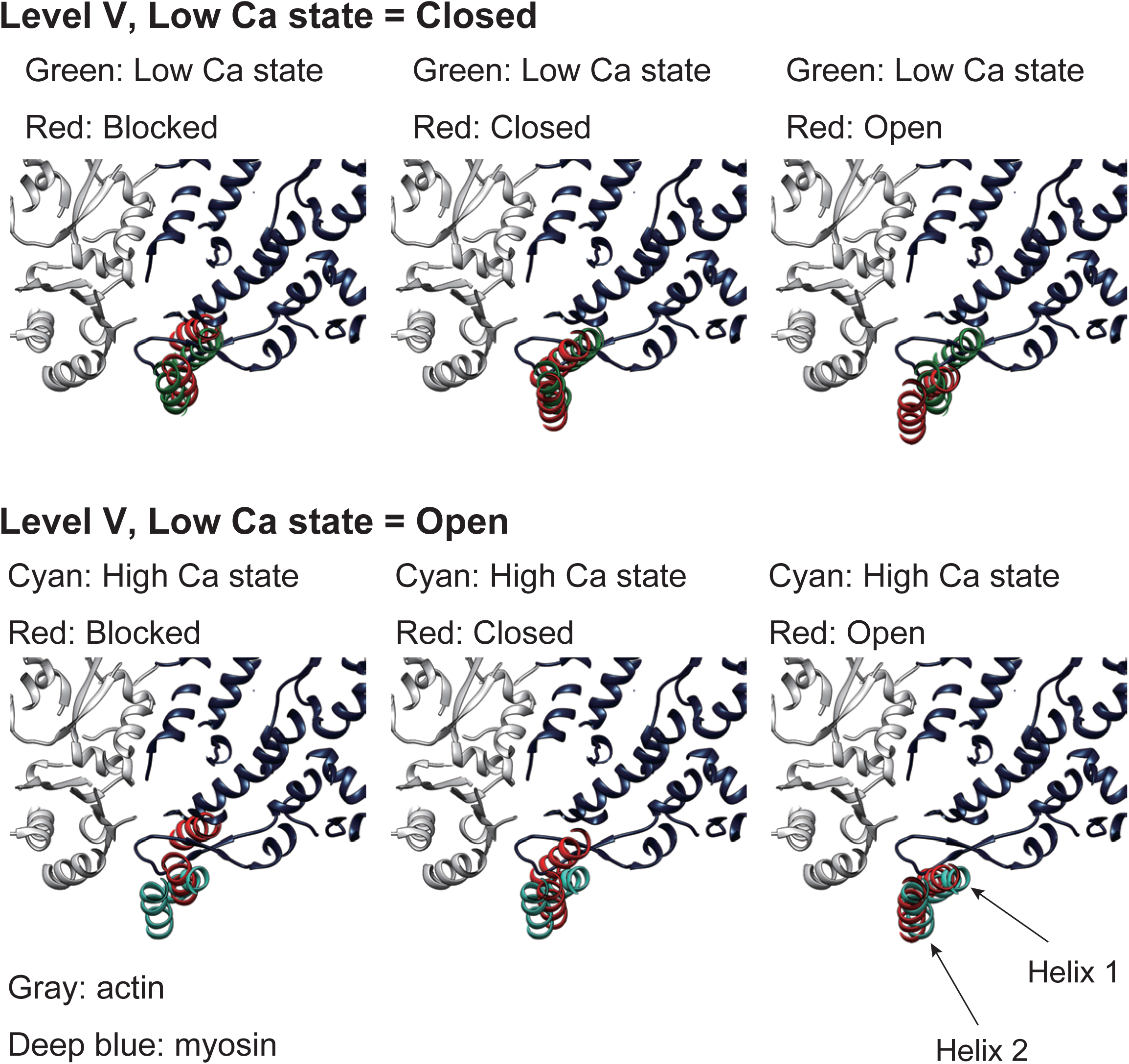
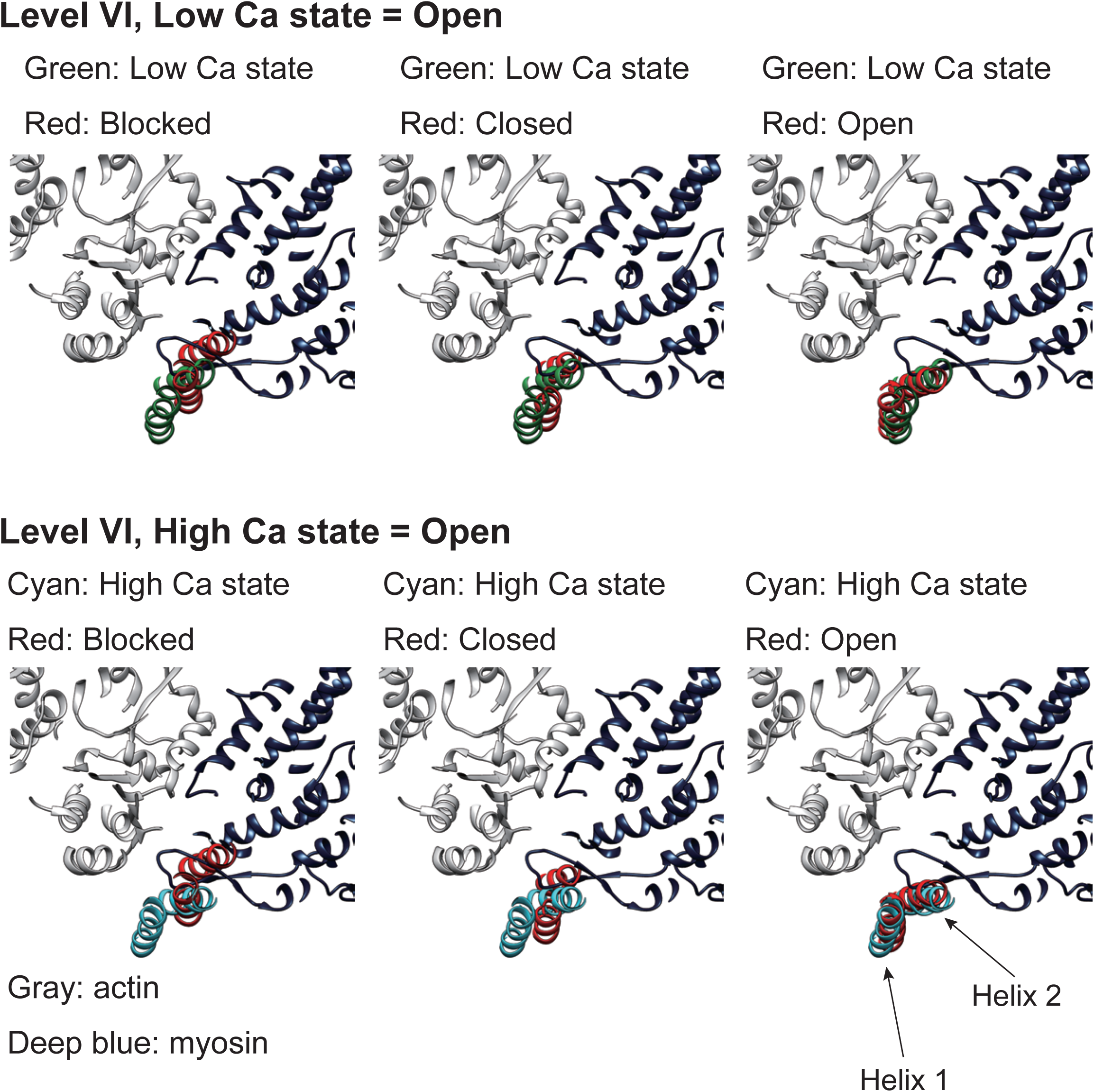
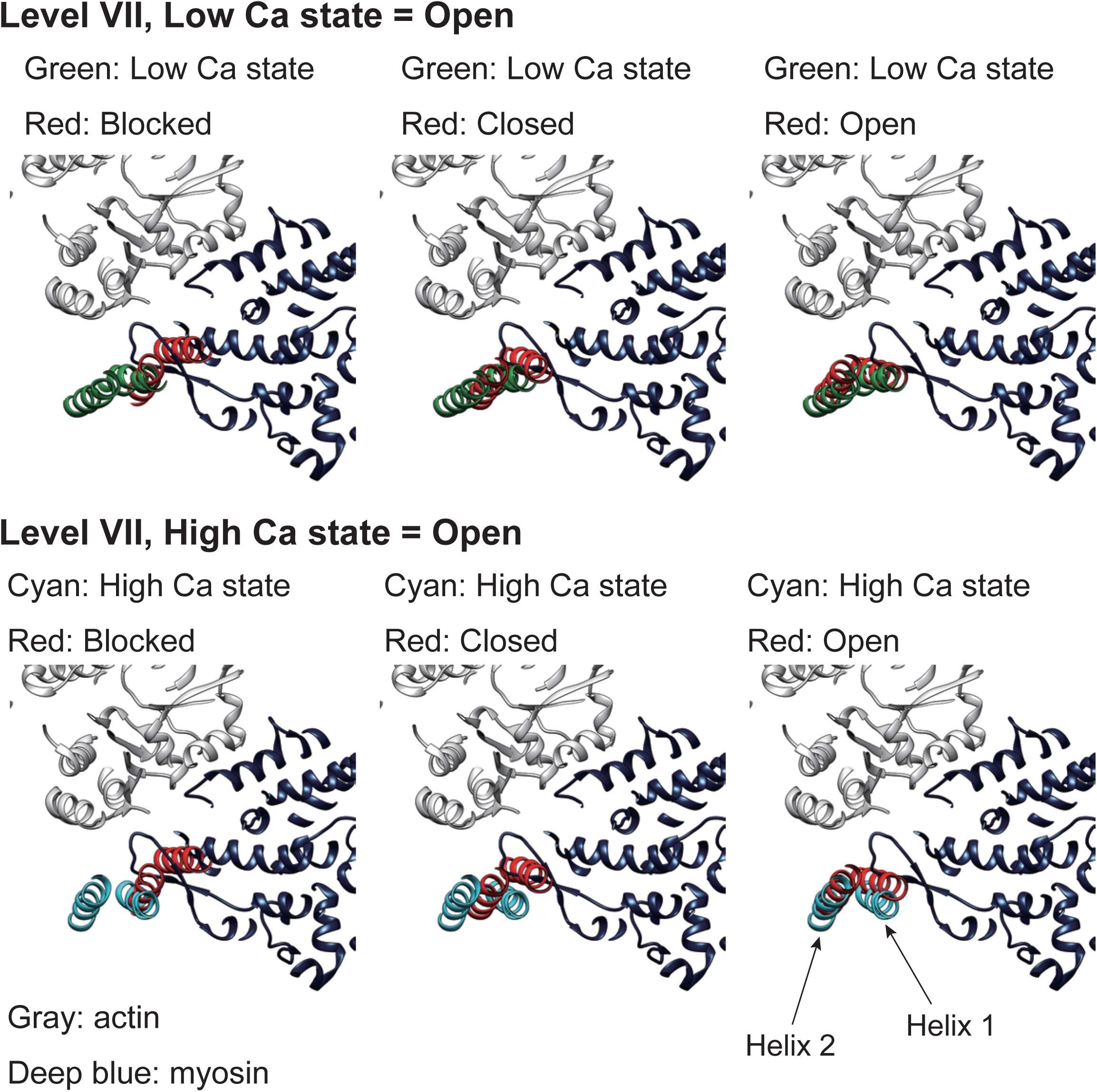
Characterization of tropomyosin positions. Cross sections showing the spatial relationships among our tropomyosin models in two calcium states (Low Ca: green; High Ca: cyan), the model of the actin-myosin complex (PDB ID: 6C1G, gray: actin; deep blue: myosin head), the models of tropomyosin in “Blocked” (PDB ID: 5NOG), “Closed” (PDB ID: 5NOL), and “Open” (PDB ID: 5NOJ) positions. Cross sections at each of the seven actin levels (I-VII) were displayed separately on each panel (A-G) (see Fig. S3A). Although myosin head cannot bind to F-actin at the level V due to the presence of troponin, myosin models were shown for convenience in comparison. Table 1 summarizes the designations of tropomyosin positions. See Fig. S3B for the surface-presentation of the sterical hindrance.

**Table 1.** Tropomyosin positions at each actin level. Tropomyosin positions were categorized based on comparison with the previously reported models (PDB 5NOG, 5NOJ, and 5NOL) (Risi et al 2017) (Fig. 4). Helix 1 is defined as the alpha-helix that faces troponin at the level V. “Blocked/Closed” and “Closed/Open” mean intermediate states between the two. In Fig. S4A, these intermediate states were counted as 0.5 (e.g. Blocked=0.5, Closed=0.5 for Blocked/Closed). Other states were counted as 1, and the percentages were calculated by dividing the sum by 14 (e.g. 8/14= 57% for the “Closed” tropomyosin in low calcium state).

| Actin levels | Low Ca state |  | High Ca state |  |
| --- | --- | --- | --- | --- |
|  | Helix 1 | Helix 2 | Helix 1 | Helix 2 |
| **I** | **Open** | **Closed?** | **Open** | **Closed?** |
| **II** | **Closed** | **Closed** | **Closed/Open** | **Closed** |
| **III** | **Closed** | **Closed** | **Closed/Open** | **Open** |
| **IV** | **Blocked/Closed** | **Blocked/Closed** | **Closed/Open** | **Open** |
| **V** | **Blocked/Closed** | **Blocked/Closed** | **Open** | **Open** |
| **VI** | **Closed/Open** | **Open** | **Open** | **Open** |
| **VII** | **Open** | **Closed/Open** | **Open** | **Open** |

|  | Low Ca state | High Ca state |
| --- | --- | --- |
| **Blocked** | **2 (14%)** | **0 (0%)** |
| **Closed** | **8 (57%)** | **3.5 (25%)** |
| **Open** | **4 (29%)** | **10.5 (75%)** |

Although many of the tropomyosin segments were categorized into either “Blocked”, “Closed”, or “Open” positions based on the comparison with the models (PDB IDs: 5NOG, 5NOJ, and 5NOL) (Risi et al 2017), some segments were in intermediate positions between the two of the three states (“Blocked/Closed” and “Closed/Open”, Table 1). At the level I, the helix 2 was in a unique position and cannot be categorized into the three states, probably because the coiled-coil structure of tropomyosin was discontinuous at the head-to-tail junction (Fig. 5, see the next section). We designated the state of helix 2 at the level I as “Closed” position for convenience in the analysis.

**Fig. 5.**
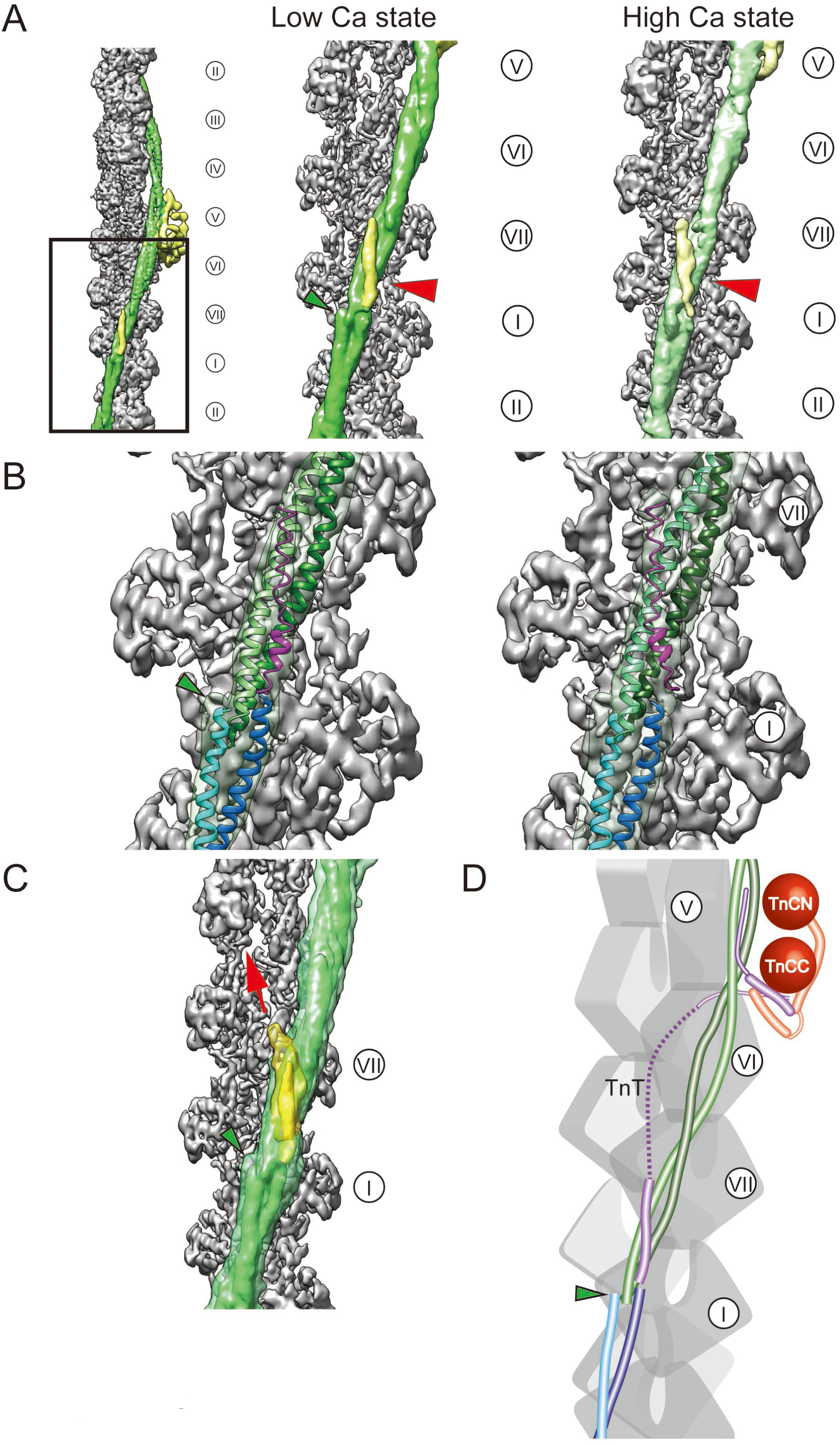
3D architecture of the head-to-tail junction of tropomyosin. (A) Thin filaments corresponding to the actin levels V-VII and I-II were reconstructed by applying an axial shift (Fig. S3A). Red arrowheads indicate the N-terminal tropomyosin-binding domain of TnT. **(B)** Crystal structure of the N-terminal domain of TnT in purple was fitted into the maps. **(C)** Lowering the threshold of the isosurface reveals that the tropomyosin-binding domain of TnT in yellow extends toward F-actin (red arrow), deviating from tropomyosin. **(D)** Schematic showing a possible conformation of TnT. The broken line in purple indicates the possible path of the middle segment of TnT. **(A-D)** The coiled-coil structure of tropomyosin appears to be disrupted at the N-terminal region, and the end region of one of two alpha-helices was visible (green arrowheads).

It appears to be contradicting that 29% of the tropomyosin segments were in “Open” position even under low calcium condition (Table 1, Fig. S4A). This is not the effect of glutaraldehyde fixation because the previous cryo-EM study of native/unfixed cardiac thin filament also demonstrates that ∼26% of the tropomyosin segments are in “Open” position under the low calcium condition (Fig. S4A) (Risi et al 2017). It is noteworthy that there is a significant sterical hindrance between the myosin head and tropomyosin in “Open” position (Fig. S3B, right), as previously shown (Risi et al 2017). Thus, these observations suggest that an “Open” tropomyosin does not fully expose the actin binding site, and a complete exposure requires a myosin binding, which displaces tropomyosin even further.

### Binding of TnT to the head-to-tail junction of tropomyosin

When we reconstructed the entire repeat of the thin filament, we noticed an extra density located at the head-to-tail junction of tropomyosin (Fig. 5A, red arrowheads). This density is likely to be the N-terminal domain (T1) of TnT, which binds to the tropomyosin junction (Narita et al., 2001; Murakami et al., 2008) (Fig. 5B, purple). The alpha-helices were discontinuous at the junction and there was an abrupt ∼90° change in the orientation of the coiled-coil (Fig. 5, green arrowheads). Although the tropomyosin binding arm of TnT has been hypothesized to run alongside tropomyosin (Gordon et al., 2000), our maps suggest that the middle segment of TnT locates away from tropomyosin (Fig. 5C), and TnT appeared to associate with tropomyosin only at the N- and C-terminal domains (Fig. 5D, purple).

The gap between the junction-associated domain of TnT and the IT arm appears to be connected by the central 59 residues of TnT (Gln182-Ser240) (Murakami et al., 2008; Takeda et al., 2003). Given that the distance between the C-terminal end of the junction-associated domain of TnT (Glu181 in PDB 2Z5H) and the N-terminal end of the IT arm (Gly241 in PDB 4Y99) was measured to be ∼109 Å in our map, part of the 59 residues of TnT are expected to form random coils because the length of a 59-residue long alpha-helix is estimated to be 88.5 Å. This calculation agrees with the previous molecular dynamics simulations (Manning et al., 2011), which shows that one half of this central linker region is predicted to form random coils.

### Actin binding sites of TnI

To identify the actin-binding domain of troponin, we subtracted the EM map of bare actin filaments (Chou and Pollard, 2019) (Fig. 6A, gray) from our thin filament maps. In both calcium states, we found a stretch of densities extending from the IT arm to Asp25 of actin at the level V (Fig. 6A, red arrowheads). These densities could be the inhibitory region of TnI, which associates tightly with the N-terminal region of actin (residues 19-44, Levine et al., 1988). Although it has been proposed that the inhibitory region (residues 137-146 of TnI) detaches from F-actin under high calcium condition, there is no direct evidence of physical dissociation of the inhibitory region from F-actin (Ishikawa and Wakabayashi, 1999). Our maps suggest that the inhibitory region of TnI stably interacts with actin and provides a supporting point for troponin. Additionally, we found a small density neighboring the N-terminal domain of actin at the level IV in low-calcium state (Fig. 6A-B, black arrowheads). This is possibly a part of the mobile domain (residues 164-210) of TnI, which binds to the intermolecular groove of F-actin (Murakami et al., 2005), but the intensity of the density was not high enough to draw a firm conclusion.

**Fig. 6.**
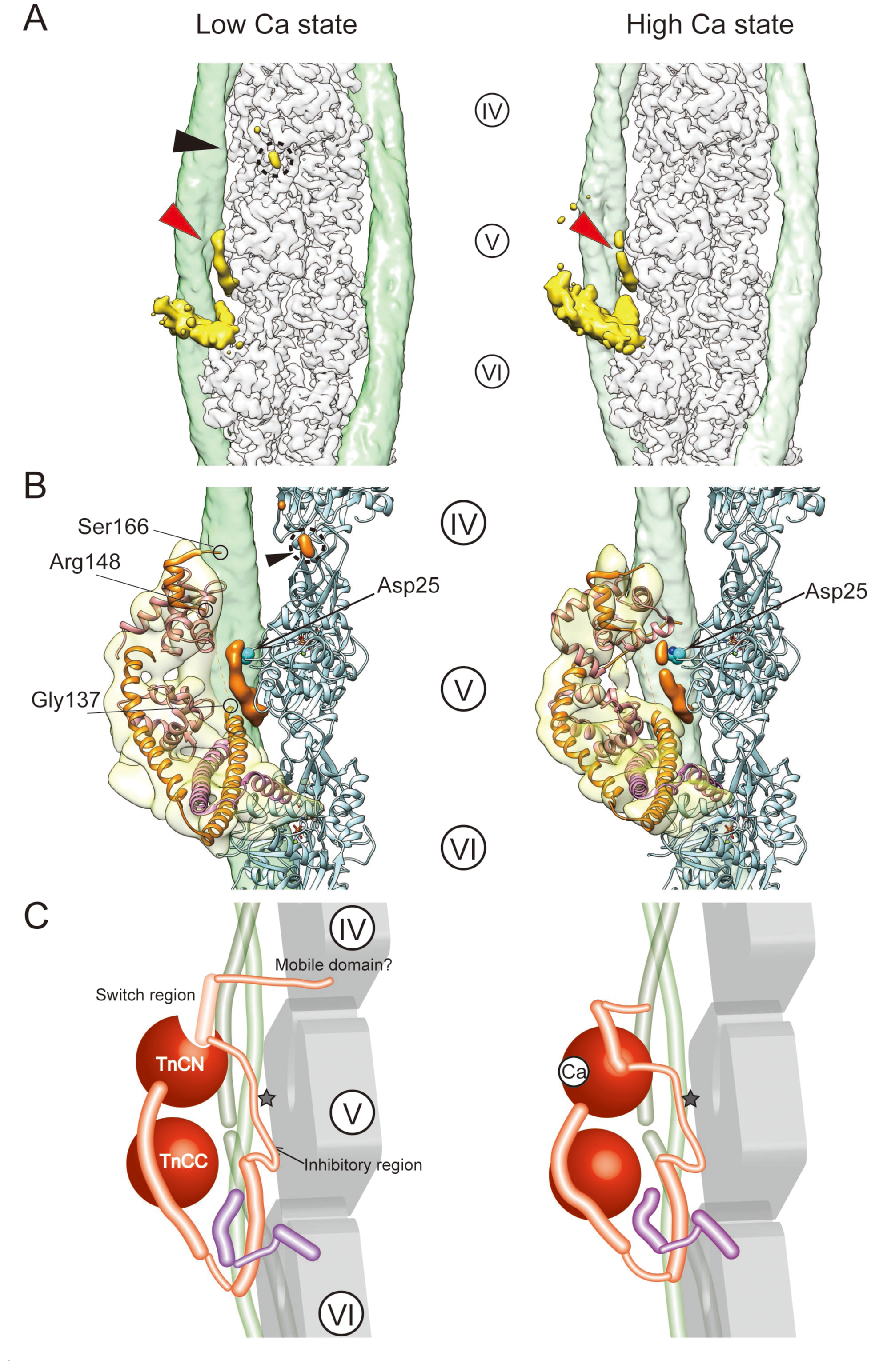
Association of TnI with F-actin. (A) Difference maps between the bare F-actin (gray) and our thin filament map. Yellow: troponin-derived difference; green: tropomyosin-derived difference. Red arrowheads: TnI inhibitory region; black arrowhead: TnI mobile domain (encircled). **(B)** Fitted crystal structures showed close association of TnI inhibitory domain with Asp25 of actin. **(B, left)** Positions of Gly137, Arg148, and Ser166 of TnI (orange) are indicated. Note that the inhibitory region (TnI 137-146) and the mobile domain (TnI 164-210) were not visualized in the crystal structure of troponin (PDB ID: 4Y99). **(C)** Schematic of the calcium-dependent conformational changes in TnI. Gray stars: Asp25 of actin.

## Discussion

### Effect of glutaraldehyde fixation

Glutaraldehyde-fixation has recently been used widely in cryo-EM, especially for challenging specimen (Stark, 2010; Adamus et al., 2019). Chemical fixation stabilizes fragile complexes and does not disrupt high resolution information at least up to 3.5 Å (Allu et al., 2019). In the case of the thin filaments, the effect of glutaraldehyde-fixation on the activity of actomyosin system has been characterized in detail (Risi et al., 2017): Fixed thin filament retains the calcium-dependent myosin-activating capability, and the calcium-induced tropomyosin shift is mostly unaffected. Differences between the native and fixed thin filament is that fixation increases the proportion of tropomyosin remaining in the blocked position under high calcium condition (Fig. S4A). Slight reduction in the myosin-activating capability and different tropomyosin behavior in fixed thin filaments may indicate that fixation reduces the mobility of tropomyosin and/or troponin. Note that we fixed the thin filament after changing the calcium concentration, thus the observed alteration in the chemical activity of the fixed thin filament does not directly represent the condition of our cryo-EM specimen. To examine the effect of glutaraldehyde on our tropomyosin structures, we compared our models with corresponding native structures (Fig. 4, Fig. S4A, and Table 1) (Risi et al., 2017). The distribution of tropomyosin positions was similar to the previous report. Based on these results of comparison, we concluded that glutaraldehyde fixation did not cause significant alterations in our structures.

### Estimation of the resolutions

It is difficult to estimate the resolution of multibody-refined maps when the resolution of each body greatly differs from one another. Thus, it is not clear whether the FSC_half-map_ curves of our troponin and tropomyosin maps (Fig. S3A) reflect their true resolutions. To make a reasonable estimation of resolution, we estimated the resolution based on the comparison between the maps and the fitted models using PHENIX validation tools (Fig. S3A, Table S2) (Adams et al., 2010). The FSC_model=0.5_ values indicate that the resolutions of the troponin and tropomyosin maps are ∼10 Å and ∼7 Å, respectively. The appearance of alpha-helices (Fig. S5B) and comparison of model-based maps filtered with various resolutions (Fig. S6) suggest that the actual resolutions of troponin maps are ∼12 Å. Our tropomyosin maps appear comparable to the 6.5 Å-map of tropomyosin (von der Ecken et al., 2015), thus the resolutions of our tropomyosin maps are expected to be similar to that of the previous map. In our actin maps, majority of the side chains were readily visible. Therefore, the FSC_half-map_-based estimation of ∼3.5 Å resolution is reliable, and the model-based validation supported this conclusion (Fig. S5A; Table S1).

### Comparison with the previous model of troponin

Our structures of troponin are different from the commonly accepted model, which assumes that the IT arm of troponin lies parallel to tropomyosin (Paul et al., 2017) (Fig. 2B, right). This “inverted” model is derived from multiple biochemical studies (Gordon et al., 2000; Perry, 1998). Therefore, it is necessary to scrutinize our model from the perspective of these previous biochemical studies.

Orientations of troponin and tropomyosin have been studied using fluorescence resonance energy transfer (FRET) (Kimura-Sakiyama et al., 2008; Miki et al., 2012). By measuring the FRET efficiency between fluorescence-labeled residues of TnT and tropomyosin, they found that the N-terminal part of the IT arm is more distant from tropomyosin than its C-terminal part (Kimura-Sakiyama et al., 2008). This observation is consistent with our model because the C-terminal end of the IT arm (Fig. S4B, Asn271 of TnT) locates close to tropomyosin, whereas its N-terminal part (Fig. S4B, Asn225 of TnT) forms the opposite end, away from tropomyosin. In the “inverted” model, however, the IT arm and tropomyosin are parallel to each other, thus the distance between the two would be nearly constant regardless of the position of the labeled residue of TnT.

Our model is also supported by the results of polarized fluorescence analysis, which shows that the helix D of TnC is approximately perpendicular to the long axis of actin (Ferguson et al., 2003) (Fig. S4C, left). In the “inverted” model, on the other hand, the helix D is almost parallel to the actin axis (Fig. S4C, right).

The interaction between tropomyosin and TnT is also important for defining the orientation of troponin. There are two tropomyosin binding sites in TnT: The N-terminal and central “T1” domain and the C-terminal “T2” domain. Because the T1 domain binds to the head-to-tail junction (Murakami et al., 2008) (Fig. 5), our conclusion with T1 agrees with the previous model. By contrast, there is a controversy on the definition of the T2 domain (Katrukha, 2013). One group claims that the T2 domain spans residues 197-239 of TnT, corresponding to part of the IT arm (Jin and Chong, 2010), whereas other groups designate the last 16 residues of TnT as T2 (Franklin et al., 2012; Tanokura et al., 1982; Morris and Lehrer, 1984). Although the former idea is consistent with the “inverted” model, we argue that the last 16 residues of TnT forms the T2/TnT hook, which strongly binds to tropomyosin as it runs between the two helices of tropomyosin (Fig. 3).

### “Flexible” tropomyosin model

The molecular mechanism of calcium-dependent myosin-activation by thin filaments depends on the structural nature of tropomyosin because one tropomyosin dimer takes a very extended coiled-coil structure, which spans seven actin monomers (Moore et al., 2016). The traditional “rigid” tropomyosin model assumes that all the seven segments/pseudo-repeats of the tropomyosin coiled-coil translocate synchronously as a single rigid body and behave like a “binary switch”: Under low calcium condition, one tropomyosin dimer blocks all the myosin binding sites on the seven actin monomers. Conversely, it exposes all the seven myosin binding sites under high calcium condition. If this model is correct, activation of myosin would not occur under low calcium condition. However, it has been shown that thin filaments enhance the phosphate release of myosin ∼8-fold even under low calcium condition (Heeley et al., 2002). Accumulating biochemical evidences indicate that tropomyosin is rather flexible, and each segment of tropomyosin can occupy different regions of actin (Maytum et al., 1999, Ishikawa and Wakabayashi, 1999). Instead of switching the positions of tropomyosin in a binary manner, calcium ions merely modify the proportions of tropomyosin segments in “Blocked”, “Closed”, and “Open” states (Fig. S4A) (Risi et al. 2017).

This “flexible” tropomyosin model agrees with our observation that the largest shift in tropomyosin occurred at the actin levels in the vicinity of troponin (Fig. 4; Table 1). This observation is reasonable because the point of action by troponin against tropomyosin locates only at the level V. Although only a subset of tropomyosin segments can be displaced by troponin, tropomyosin shifts could be amplified and propagated along the long axis of F-actin via cooperative binding of myosin (Maytum et al., 1999; Tobacman and Butters, 2000).

## Materials and Methods

### Isolation of thin filaments

Native murine cardiac thin filaments were isolated along with thick filaments from ventricular myocardium of six adult ICR mice (∼24 weeks old). Mice were anesthetized with intra-abdominal injection of medetomidine, midazolam, and butorphanol. Once anesthetized, mice were killed by injection of lethal dose (80 mg/kg) of pentobarbital. These procedures were approved by the University of Yamanashi Animal Experiment Regulation Committee. Immediately after euthanasia, the heart was excised and placed in a hypotonic buffer (Hepes 30 mM, pH 7.2) supplemented with protease inhibitor cocktail (Nacalai tesque, Kyoto, Japan), and atria, aorta, and pericardium were carefully removed. The remaining ventricular myocardium were gently homogenized with a Dounce glass homogenizer (Wheaton, Millville, NJ). The homogenized tissue was washed three times with a hypotonic HK buffer (Hepes 30mM, pH 7.2, 60 mM KCl, 2 mM MgCl2, protease inhibitor cocktail) and cleared by centrifugation at 750 ×g for 15 min at 4 ℃. The washed tissue was incubated in HK buffer plus 1% Triton X-100 for 15 min at 4℃. The demembranated tissue was washed again three times with HK buffer and was subsequently incubated in HK buffer plus 2 mM ATP and 1 mM EGTA. The ATP-treated tissue was disintegrated by passing through a 23G needle five times and the debris was removed by centrifugation at 10,000 ×g for 10 min at 4 ℃. The supernatant was further processed for cryo-EM.

### Sample preparation for cryo-EM

For high-calcium state, CaCl2 (final 2 mM) and sodium orthovanadate (final 1 mM) were added to the supernatant and incubated for 15 min at 4 ℃. Sodium orthovanadate was added to reduce binding of myosin to the thin filaments. For low-calcium state, only sodium orthovanadate (final 1 mM) was added to the supernatant. After changing the calcium concentration, the sample was fixed with 0.25% glutaraldehyde for 5 min at 20 ℃, according the previous study (Risi et al., 2017), and subsequently quenched with Tris-HCl (final 50 mM, pH 7.2) for another 5 min at 20℃. The protein concentration of the sample was adjusted to 0.1 mg/ml and 3 µl of the sample was mounted on freshly glow-discharged holey carbon grids, Quantifoil R1.2/1.3 Cu 200 mesh (Quantifoil Micro Tools GmbH, Großlöbichau, Germany), blotted for 10 sec at 4℃ under 99% humidity, and plunge frozen in liquid ethane using Vitrobot Mark IV (Thermo Fisher Scientific, Waltham, MA). Grid quality was examined using JEM-2100F (JEOL, Tokyo, Japan) at University of Yamanashi. Grids frozen under optimal conditions were then used in the recording session.

### Image acquisition

Images were recorded using a Titan Krios G3i microscope at University of Tokyo (Thermo Fisher Scientific) at 300 keV equipped with a VPP, a Gatan Quantum-LS Energy Filter (Gatan, Pleasanton, CA) with a slit width of 20 eV, and a Gatan K3 Summit direct electron detector in the electron counting mode. The nominal magnification was set to 81,000× with a physical pixel size of 1.07 Å/pixel. Each movie was recorded for 5.6 sec with a total dose of 60 electrons/Å2 and subdivided into 50 frames. Movies were acquired using the SerialEM software (Mastronarde, 2005) and the target defocus was set to 0.4 µm. The VPP was advanced to a new position every 15 min. The image acquisition was significantly accelerated (∼8,800 movies in 48 hours) by a multiple exposure scheme utilizing the beam-tilt, which acquires movies from nine different holes every stage shift (Zivanov et al., 2018).

### Data processing

Movies were subjected to beam-induced motion correction using MotionCor2 (Zheng et al., 2017), and the contrast transfer function (CTF) parameters were estimated using Gctf v1.18 (Zhang, 2016). Initially, thin filaments were manually selected, and ∼5,000 segments were subjected to 2D classification for generating reference class averages for auto particle picking. Using the reference-based autopick function of Relion-3 (Zivanov et al., 2018), 3× binned 86 × 86 pixel segments (3.21 Å/pixel) were extracted from ∼8,300 micrographs.

Since auto-pick subroutine failed to distinguish thin filaments from thick filaments, we performed five rounds of 2D classification to eliminate thick filaments and debris. Then, unbinned 264 × 264 pixel segments (1.07 Å/pixel) of the thin filaments were re-extracted and were subjected to three more rounds of 2D classification. Selected segments were further subjected to 3D refinement, CTF refinement, beam-tilt refinement, and Bayesian polishing cycles. For evaluate the quality of the datasets, we reconstructed the F-actin part imposing helical symmetry with helical twist of −166.6° and helical rise of 27.7 Å using a mask excluding troponin and tropomyosin (Table S1, F-actin (3D auto-refine, helical symmetry imposed)). These symmetry-imposed maps were not used for the main analysis of this study, and no helical symmetry was imposed for 3D refinement of the whole thin filament and for the subsequent multibody refinements.

To classify the segments according to the presence and position of troponin, align-free 3D classification was performed. Segments were classified according to the position of troponin along the filaments (Fig. S1 and S2). Segments without troponin densities were discarded and the remaining segments were re-extracted with axial-offsets to bring the troponin density to the box center. After duplicates removal and several rounds of 3D classification, troponin densities were converged to a single class, which had a troponin density on each side of the filament.

The two troponin densities were expected to be combined by applying local P2_1_ symmetry with −166.6° twist and 27.7 Å rise. However, we found that positions of the two troponin densities did not conform to this local symmetry due to high structural heterogeneity. Thus, we re-extracted two sets of segments from a single set of the refined segments; one set was extracted without axial offsets, and the other was extracted with a 27.7-Å axial offset. By combining these two sets of segments, we merged the troponin densities on both sides of the thin filaments in a single process of 3D auto-refinement. After merging the two sets of the segments, the refined 3D structure had a strong troponin density on one side, while having weak and noisy troponin densities on the other side. These noisy troponin densities appeared because the 3D refinement was applied only to one of the troponin pair, and the other troponin was blurred out. Therefore, we ignored these noisy troponin densities on the back side of the filament in the subsequent data processing.

After removal of duplicates, segments were re-extracted and subjected to 3D auto-refinement without applying helical symmetry. We created a loose mask enclosing the troponin region and performed a focused 3D classification with image alignment. The major class, which showed structural details the most, were selected and further subjected to a focused 3D classification without image alignment. The remaining minor classes were in very low resolution or part of the troponin L-shape was truncated, suggesting that they were poorly aligned probably due to low-contrast. When the number of classes was increased from four to seven, more minor low-resolution/truncated classes appeared, whereas the major high-resolution class was mostly unaffected. Truncations in the structure of minor classes usually occur either within the TnC part or the IT arm, suggesting that the possible hinge between the subdomains (Fig. 2A) is the major cause of structural heterogeneity in troponin. A large regularization parameter of T=20 was used for enhancing the high-resolution information. Changing the T values between 10 to 20 did not change the result. To visualize the spatial relationship among actin, tropomyosin, and troponin, a three-body refinement was conducted using three corresponding masks (Fig. S1 and S2) (Nakane et al., 2018). Since identification of the inter-body boundary between tropomyosin and troponin was crucial in this study, we created a loose mask with 10-pixel extension and 10-pixel soft-edge for the troponin body. There was a large overlapped region between the masks of troponin and tropomyosin, avoiding arbitrary placement of the inter-body boundary. In contrast, the mask for F-actin did not overlap neither with masks for tropomyosin or troponin because even a small portion of actin densities included in tropomyosin or troponin mask could cause biased alignment toward F-actin, messing up tropomyosin or troponin bodies. Since the result of the multibody refinement cannot be used for the analysis of the actin-troponin interface, the actin-binding domain of TnI (Fig. 6) was based on the result of the 3D refinement of the whole thin filament (Fig. S1-2, upper right; Table S2, EMD-0807, 0808), and we did not use the results of the multibody refinement.

Note that the maps generated by multibody refinement represent the median position of the flexible body (Nakane et al., 2018). As maps were generated separately for each body, we prepared figures by reassembling these bodies into a single composite map.

For reconstruction of the axially-shifted regions of the filament (Fig. S3A), segments were re-extracted with axial offset either of −104 pixels, −52 pixels, or +104 pixels. Segments were similarly refined using 3D auto-refine and multibody refinements.

It is known that TnC, TnI, and the C-terminal domain of TnT constitutes the “head” or “core” of troponin (Takeda et al., 2003). The remaining N-terminal domain of TnT is proposed to run along the tropomyosin until it reaches the junction of two adjacent tropomyosin molecules, which is called the “head-to-tail junction” (Murakami et al., 2008). We also tried to include both the troponin head and the head-to-tail junction within one box by enlarging the box size two-fold. However, due to the curvature of the filaments, only the central part of the filament was analyzable and resolution deteriorated toward the edges of the box.

### Model building

For model building of actin, ADP-actin model (PDB ID: 6DJO) (Chou and Pollard, 2019) was fitted to our thin filament density maps using PHENIX real-space refine subroutine (Word et al., 1999; Afonine et al., 2018; Adams et al., 2010). As for model building of troponin, the crystal structure of the human cardiac troponin core (PDB ID: 4Y99), an improved version of PDBID: 1J1E (Takeda et al., 2003), cannot be directly fitted into our maps. Thus, we split the crystal structure into several domains, and manually fitted them into our maps using UCSF Chimera (Pettersen et al., 2004). The roughly fitted models were further aligned using PHENIX real-space refinement. As the 4Y99 structure is calcium-saturated form, we used the NMR structure of calcium-free turkey skeletal TnC N-terminal domain (PDBID: 1TRF) (Findlay et al., 1994) for fitting the TnCN lobe in low-calcium state. For building the tropomyosin model, we fitted the alpha-helix models of F-actin-tropomyosin complex (PDBID: 3J8A) (von der Ecken et al., 2015) or the crystal structure of boar tropomyosin (PDBID: 1C1G) (Whitby and Phillips, 2000). The model of the actin-myosin complex (PDBID: 6C1G) (Mentes et al., 2018) was fitted using UCSF Chimera to visualize the collision between tropomyosin and the myosin head. To identify the actin-binding domains of TnI, we subtracted the ADP-actin map (EMD-7938) (Chou and Pollard, 2019) from our thin filament maps in low- and high-calcium states. The resulting difference maps were de-noised by applying 3×3×3 median filtering. For comparison of the cross-correlation values between the troponin map and the 4Y99-based model in different orientations (Fig. 2A), we first manually fitted 4Y99 model into the troponin map in high calcium state using UCSF Chimera. Then, the roughly-aligned models were further aligned using PHENIX real space refinement tool and the cross-correlation values were calculated using the comprehensive validation tool.

The statistics of the 3D reconstruction and the model validation are summarized in Table S1.

## Supplementary Materials

Movie S1. Movie presentation of the thin filament structure in low-calcium state.

Movie S2. Movie presentation of the thin filament structure in high-calcium state.

## Abbreviations

VPP: Volta phase plate
TnC: troponin C
TnT: troponin T
TnI: troponin I
IT arm: TnI-TnT arm
FSC: Fourier shell correlation

## Acknowledgments

**General**: This research is partially supported by Platform Project for Supporting Drug Discovery and Life Science Research (Basis for Supporting Innovative Drug Discovery and Life Science Research (BINDS)) from Japan Agency for Medical Research and Development (AMED) under Grant Number JP19am0101115. Computational resource of SGI Rackable C1102-GP8 (Reedbush-U/H/L) was awarded by “Large-scale HPC Challenge” Project, Information Technology Center, the University of Tokyo. We thank Dr. Akihiro Narita (Nagoya University) and Dr. Takanori Nakane (MRC Laboratory of Molecular Biology) for helpful discussions.

## Author contributions

T.O. designed the research; T.O., H.Y., and T.W. analyzed data and wrote manuscript.

## Data and materials availability

Deposited maps and models are summarized in Table S2.

## Supplemental Figure Legends

**Fig. S1.**
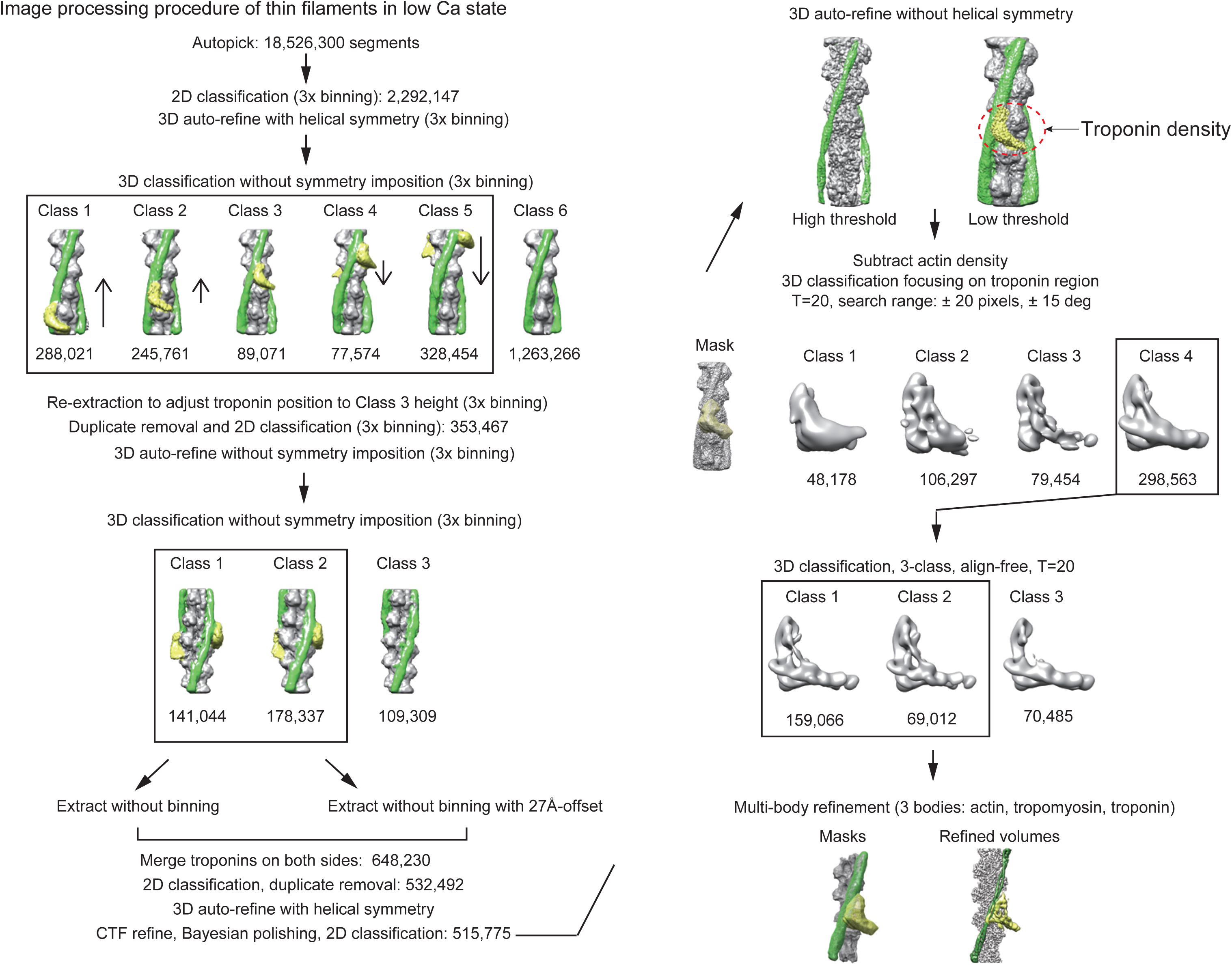
Procedure of the image analysis of the cardiac thin filaments in low-calcium state. Red broken circle (upper right) indicates the noisy troponin densities, which could not be aligned by conventional 3D refinements of the whole thin filament. The numbers indicate the number of segments used for the refinement or belonging to the indicated class.

**Fig. S2.**
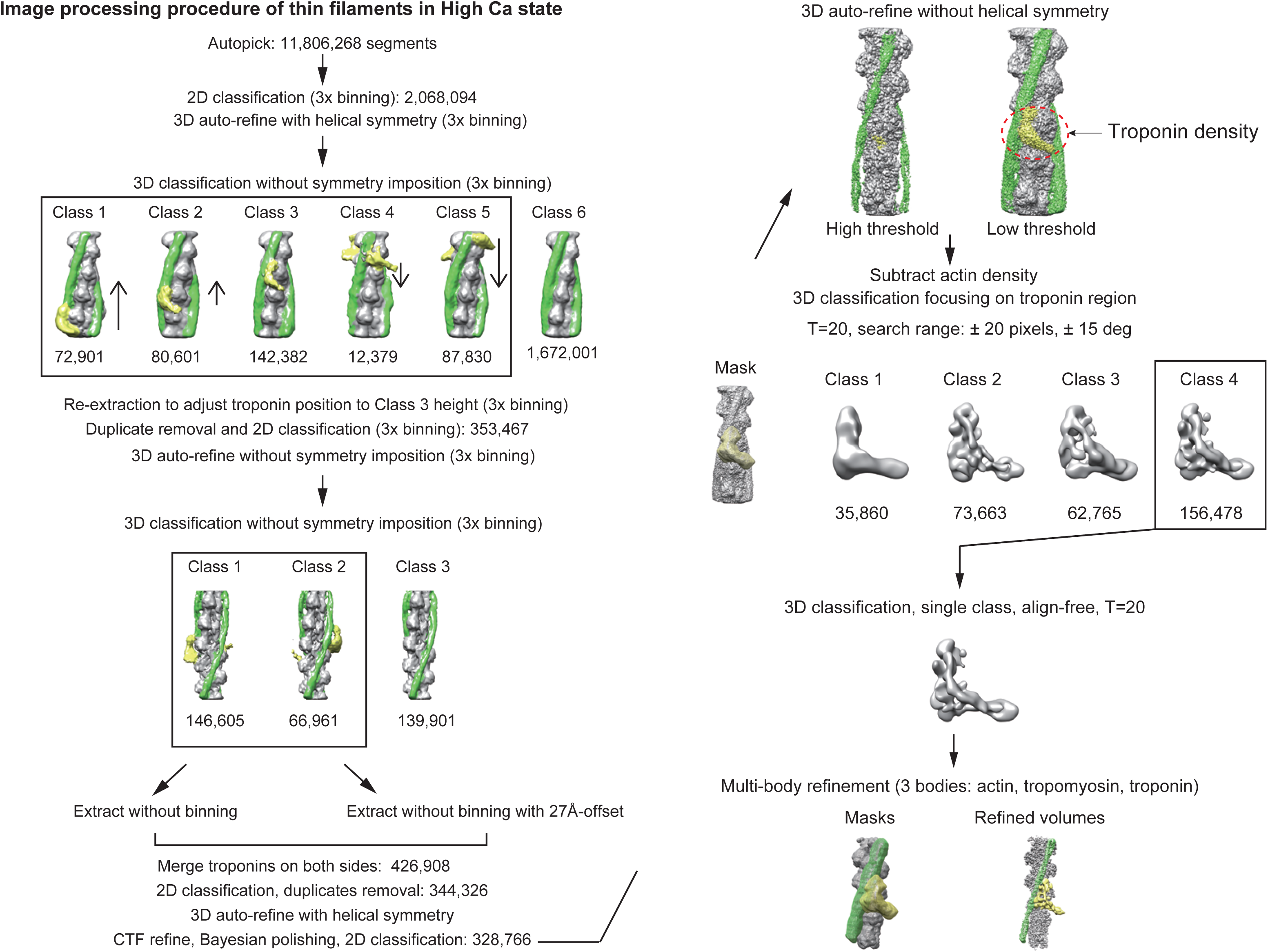
Procedure of the image analysis of the cardiac thin filaments in high-calcium state. Red broken circle (upper right) indicates the noisy troponin densities, which could not be aligned by conventional 3D refinements of the whole thin filament. The numbers indicate the number of segments used for the refinement or belonging to the indicated class.

**Fig. S3.**
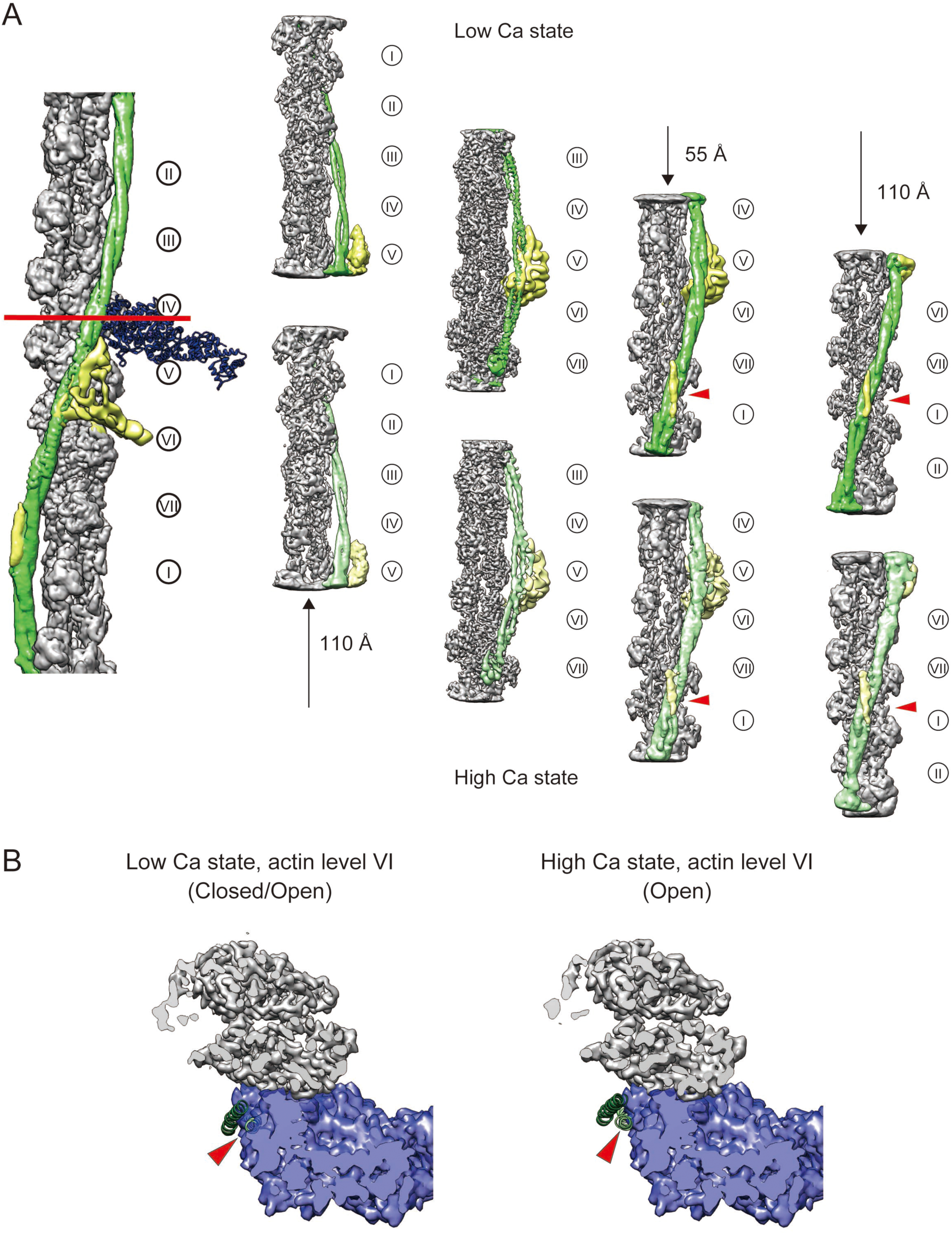
Reconstruction of the entire seven levels of the thin filament. (A) Axial offsets of +110 Å, −55 Å, and −110 Å were applied to the original segments, and the re-extracted segments were further refined using 3D auto-refine and multibody refinement. **(B)** Visualization of the sterical hindrances at the actin level VI. Even in “Open” position, there is a significant sterical hinderance between tropomyosin and the myosin head (red arrowhead, right).

**Fig. S4.**
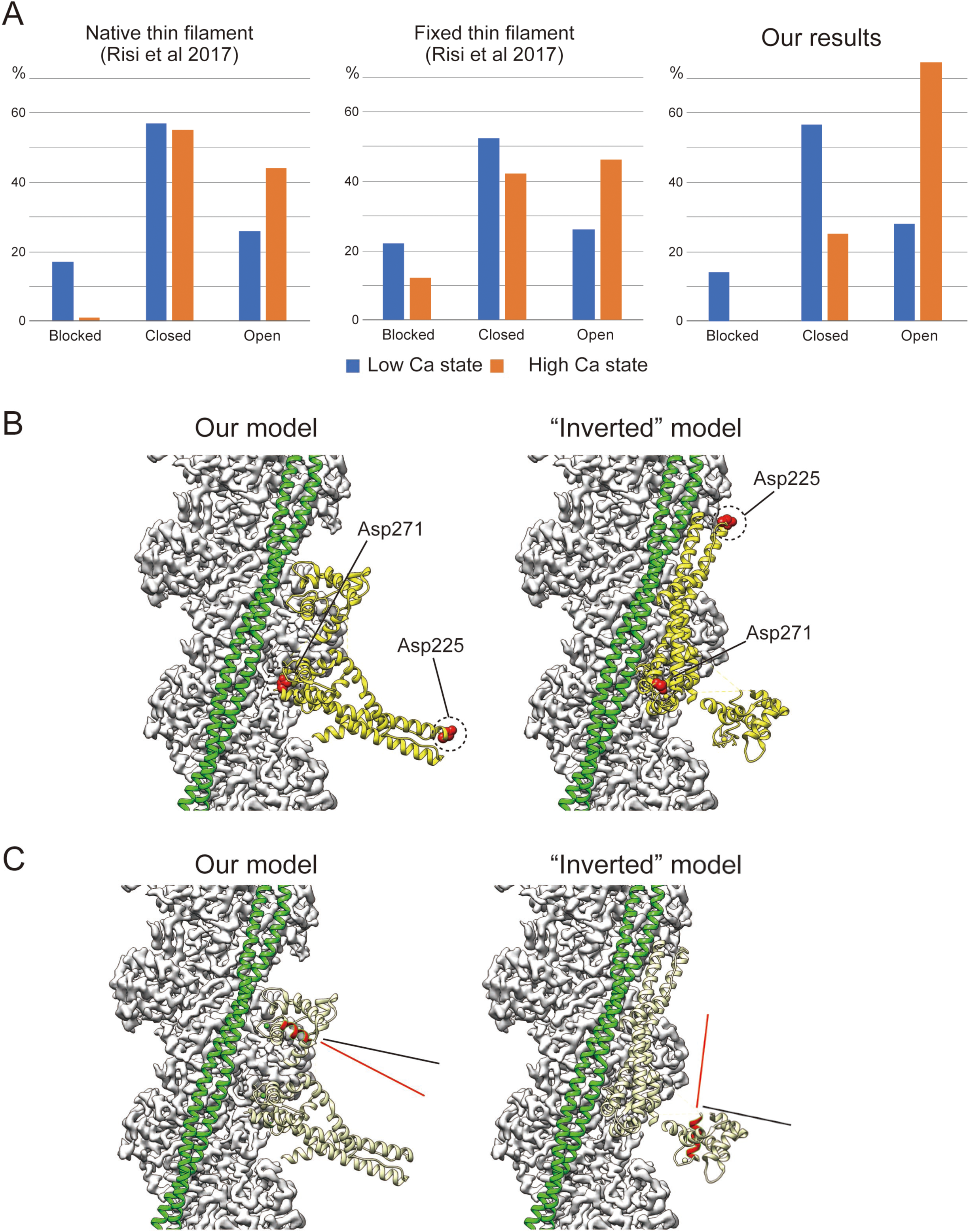
Comparison with previous reports. **(A)** Proportions of tropomyosin in each of the three states. The left and middle graphs were adopted from Fig. 1B of Risi et al 2017. See Table 1 for the calculation of percentages based on the comparison in Fig. 4. **(B)** Distance between TnT and tropomyosin. In our model, Asn225 is greatly distant from tropomyosin compared with Asn271. In the classic inverted model, by contrast, the distance between TnT and tropomyosin is essentially constant. **(C)** Orientation of the helix D (red, residues 74-83 of TnC) relative to the long axis of F-actin. According to the previous study (Ferguson et al., 2003), the angle between the helix D and the actin axis is ∼78 °. In our model, the angle is ∼62 °. In the inverted model, by contrast, the angle is ∼6 °. Red and black lines indicate the differences in the angles between the model (red) and the previous study (black, 78 °).

**Fig. S5.**
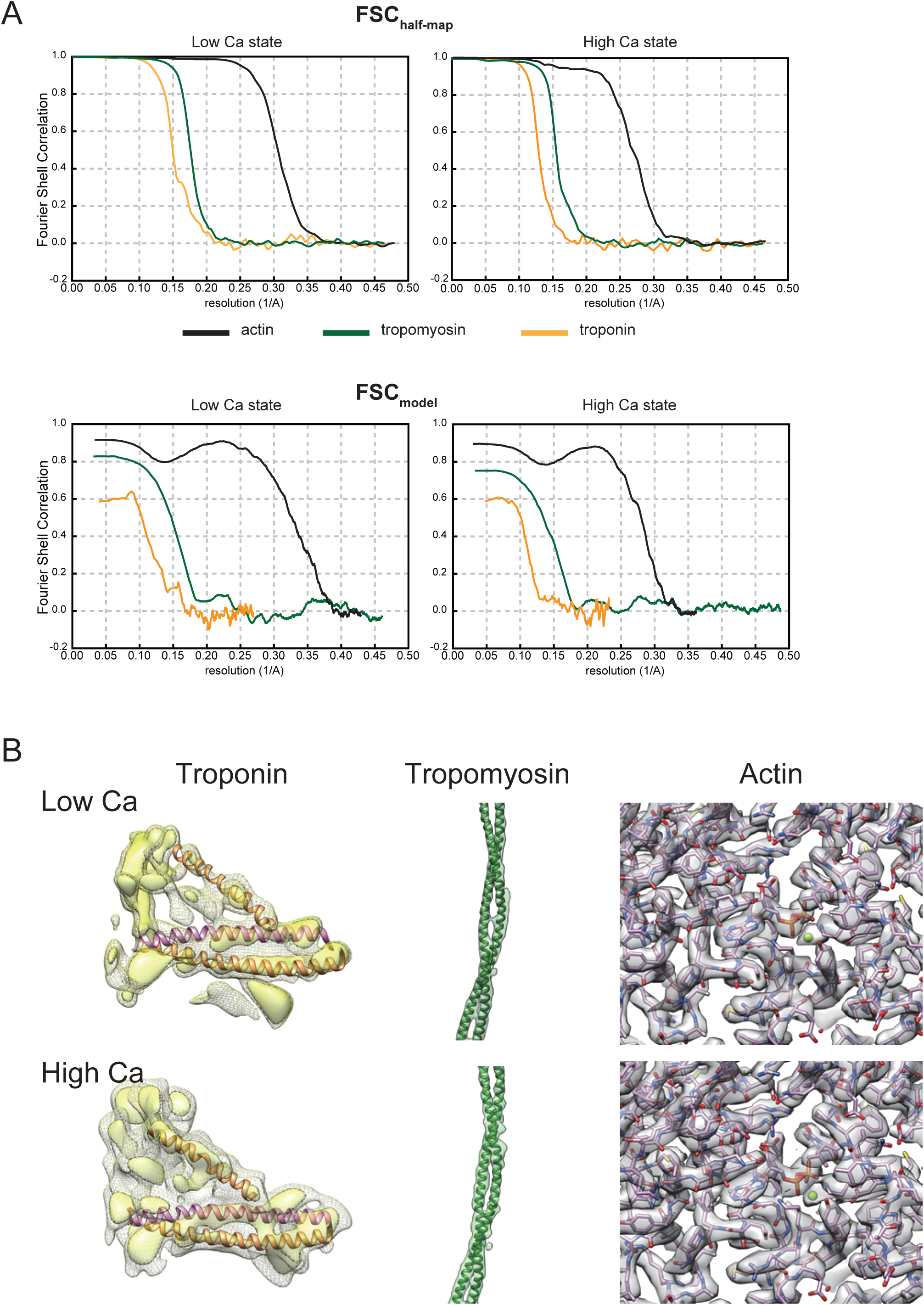
Estimation of resolution. (A) Fourier shell correlation (FSC) curves. FSC_half-map_ curves were calculated using Relion postprocess tool. FSC_model_ curves were calculated using PHENIX comprehensive validation tool. Statistics of the resolution estimation were summarized in Table S2. **(B)** The appearance of alpha-helices and side chains in the maps of multibody refinement were shown. The alpha-helices of the IT arm and the TnI brace were visible in troponin maps. The alpha-helices of the coiled-coil were well-separated in the tropomyosin maps. Side chains were well-visualized in F-actin maps.

**Fig. S6.**
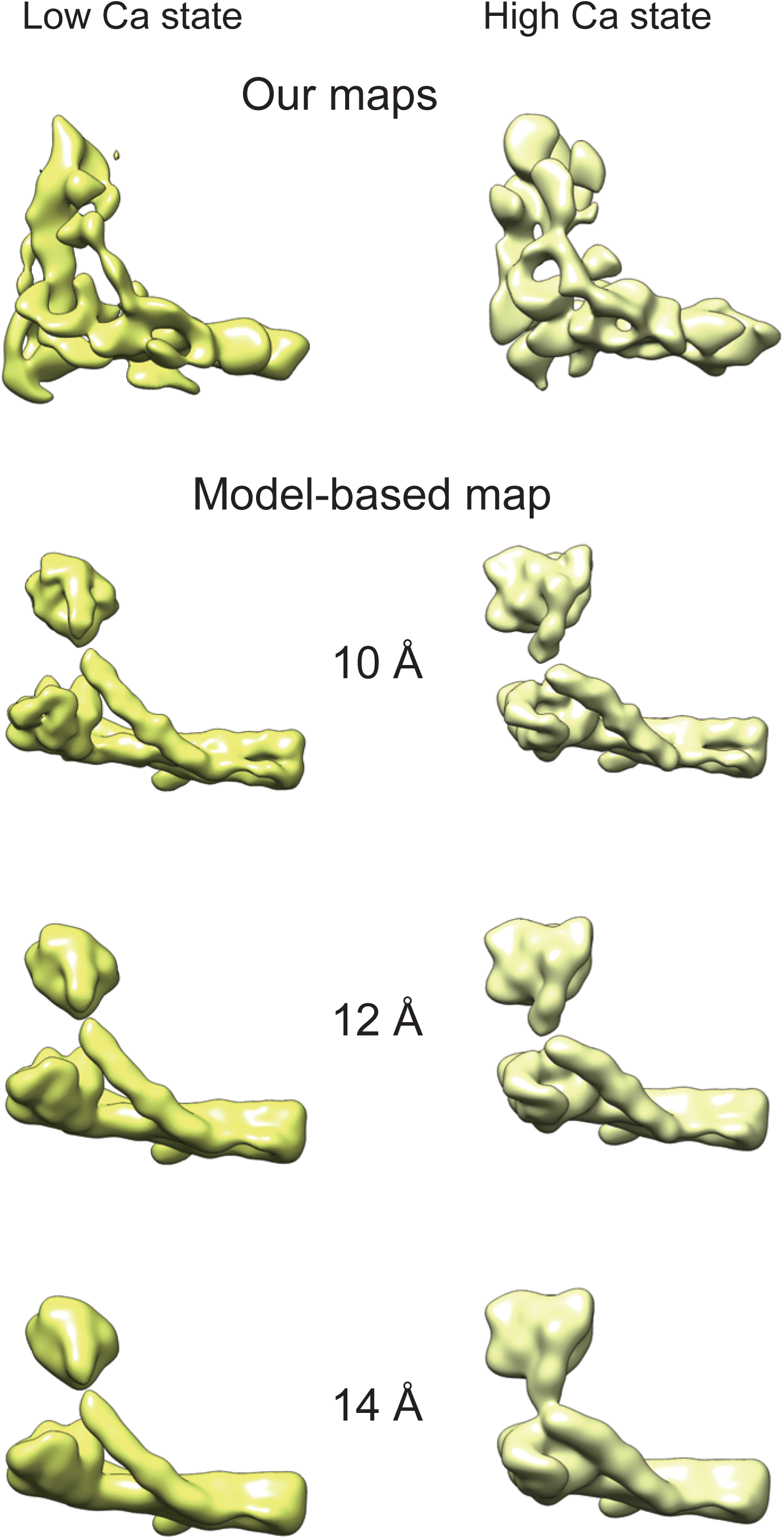
Comparison with model-based maps. Our troponin maps were compared with simulated maps derived from the fitted models. Simulated maps were low-pass filtered with indicated resolutions. As the crystal structure PDB ID: 4Y99 does not include large portions of TnI and TnT, the simulated maps were smaller than our maps. Moreover, a crystal structure of troponin in low calcium state has not been available yet, thus there is a great difference in the TnC part between our map and the model-based map. It is likely that the structure of TnC in low calcium state greatly differs from the crystal structure in high calcium state.

**Table S1.**
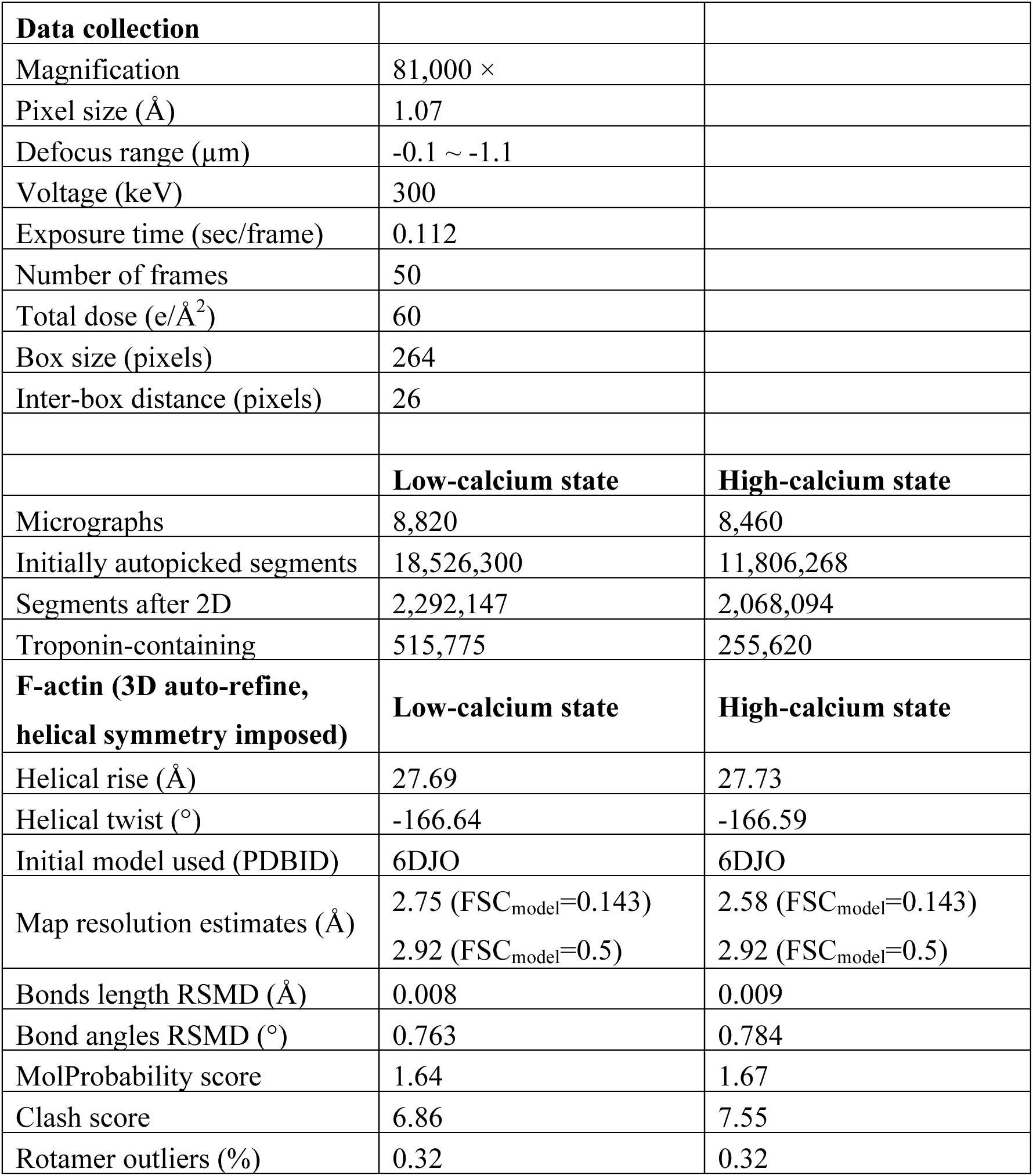
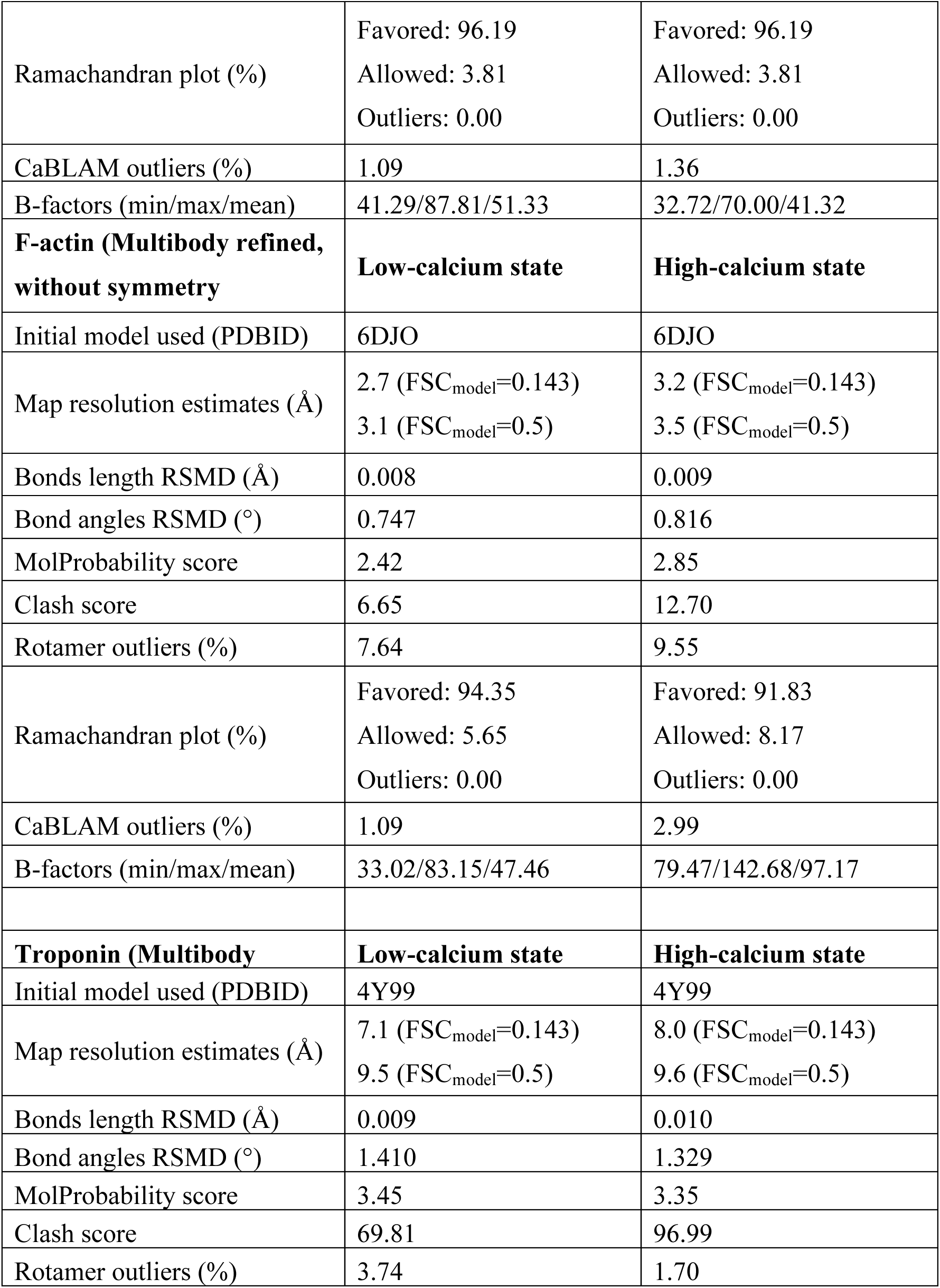
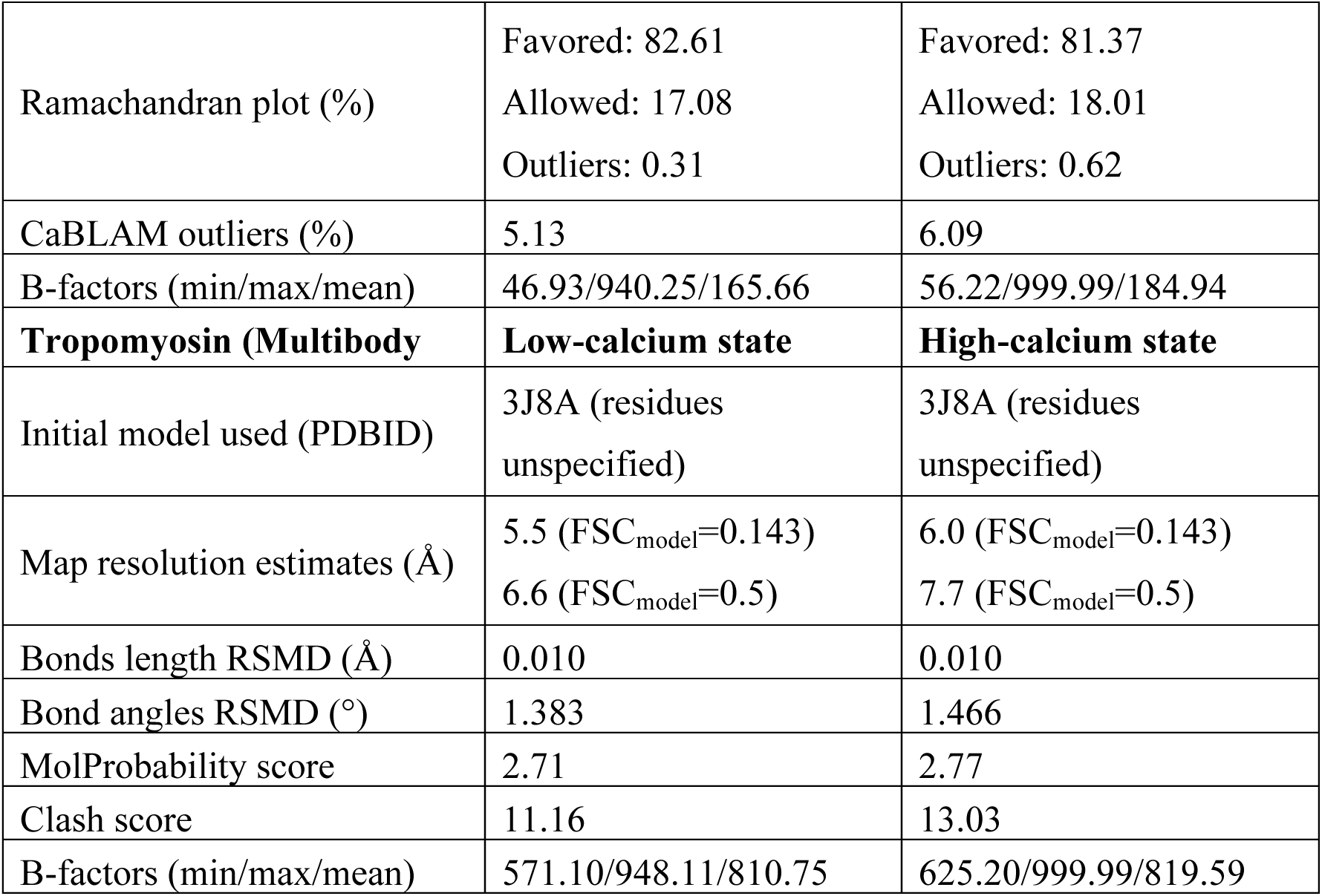
Cryo-EM structure determination and model validation statistics.

**Table S2.** Summary of deposited maps and models.

|  | EMDB ID | PDB ID |
| --- | --- | --- |
| Main structures |  |  |
| F-actin in low Ca state | 0711 | 6KLL |
| F-actin in high Ca state | 0712 | 6KLN |
| Tropomyosin in low Ca state | 0714 | 6KLP |
| Tropomyosin in high Ca state | 0715 | 6KLQ |
| Troponin in low Ca state | 0717 | 6KLT |
| Troponin in high Ca state | 0718 | 6KLU |
| Axially-offset structures |  |  |
| F-actin in low Ca state<br>actin levels II-IV | 0796 |  |
| Tropomyosin in low Ca state<br>actin levels II-IV | 0797 |  |
| F-actin in low Ca state<br>actin levels VI-VII, I | 0798 |  |
| Tropomyosin in low Ca state<br>actin levels VI-VII, I | 0799 |  |
| F-actin in high Ca state<br>actin levels II-IV | 0802 |  |
| Tropomyosin in high Ca state<br>actin levels II-IV | 0804 |  |
| F-actin in high Ca state<br>actin levels VI-VII, I | 0805 |  |
| Tropomyosin in high Ca state<br>actin levels VI-VII, I | 0806 |  |
| Non-multibody refined maps |  |  |
| Thin filament in low Ca state | 0807 |  |
| Thin filament in high Ca state | 0808 |  |

